# Single-cell genomics reveals genomic and functional differentiation between freshwater and saline *Achromatium* populations

**DOI:** 10.64898/2026.11.16.767035

**Authors:** Pierre H. Remy, Danny Ionescu, Olivier Gros, Sophie-Luise Heidig, Tanja Woyke, George Schaible, Jean-Marie Volland

**Affiliations:** Institut de Systématique, Evolution, Biodiversité, Université des Antilles, Muséum National d’Histoire Naturelle, CNRS, Sorbonne Université, EPHE, Campus de Fouillole, 97110 Pointe-à-Pitre, Guadeloupe, France; Department of Environmental Microbiomics, Technische Universität Berlin, Berlin, Germany; Interuniversity Institute for Bioinformatics in Brussels IB2, ULB-VUB, Brussels, Belgium; Department of Molecular, Cellular, and Developmental Biology, University of California, Santa Barbara, Santa Barbara, CA, United States; Department of Energy Joint Genome Institute, Lawrence Berkeley National Laboratory, Berkeley, CA, United States

**Author notes:** **Correspondence:** Jean-Marie Volland.

**Keywords:** giant sulfur-oxidizing bacteria, Caribbean mangroves, ultrastructure, single-cell genomics, intracellular diversity, comparative functional genomics

## Abstract

Giant sulfur bacteria of the genus *Achromatium* reportedly contain extensive genomic diversity within individual cells. Most genomic data (35 assemblies) come from freshwater populations obtained from Lake Stechlin, Germany, alongside four single-cell genomes from a salt marsh and saline mineral springs. Here, we characterize marine *Achromatium* from mangrove sediments in Guadeloupe and compare them with freshwater and saline references. Microscopy, elemental mapping, and Raman spectroscopy revealed calcium-rich inclusions, and intracellular elemental sulfur as previously described for this genus. Scanning electron microscopy revealed two cell-size morphotypes. We generated eleven single-cell, four mixed-cell, and two metagenomic binned assemblies from Guadeloupe mangrove. Divergent copies of conserved marker genes revealed extensive intracellular genomic diversity in individual cells of both morphotypes. Multigene phylogenies provided separation between freshwater references and most genomes from saline environments, including the Guadeloupe population. Gene presence-absence analyses further revealed differences in functional potential between Guadeloupe and Stechlin cells across five functional annotation categories including PFAM, KEGG Orthologs, and CAZymes. Differences in genes associated with osmotic regulation, ion transport, carbohydrate degradation, and secretion systems suggest variation in functional potential related to environmental adaptation, interactions and resource utilization. No significant differences in functional potential between Guadeloupe morphotypes were found. Together, these marine genomes reveal genomic and functional differentiation from freshwater populations and suggest that intracellular diversity does not erase freshwater/saline population boundaries.

## Introduction

The *Achromatium* genus contains the largest known freshwater bacteria and is one of the few that can be identified from their morphology alone. They comprise uncultured giant sulfur bacteria historically grouped under *Achromatium oxaliferum*, described in 1893 [1, 2]. Cells can exceed 100 µm in length and contain abundant calcium carbonate-rich bodies and sulfur inclusions [2–5]. The genus belongs to the *Chromatiaceae* within the *Gammaproteobacteria* and *Pseudomonadota* [6]. Populations occupy oxic–anoxic interfaces in marine and freshwater sediments, where sulfur oxidation and intracellular sulfur storage accompany versatile carbon metabolism [2, 5, 7, 8]. Microautoradiography documented assimilation of bicarbonate, acetate, and complex organic substrates by different populations or subpopulations [9–11]. Recent metagenomic [8] and proteomic [12] studies provided support for two carbon-fixation pathways and reported the expression of temperature-associated protein variants in freshwater cells. The contribution of nitrate to *Achromatium* metabolism remains unresolved, yet high nitrate concentrations elicit a negative chemotactic response [13].

The nature and localization of carbonate inclusions have also been assessed. Cellular calcium/sulfur ratios vary with depth and diel cycle in salt-marsh populations [14]. These calcium carbonate bodies likely occupy periplasmic pockets formed by cytoplasmic-membrane invaginations [15]. Raman microspectroscopy identified amorphous calcium carbonate rather than the crystalline calcite as was assumed in much of the earlier literature [16]. These findings make combined structural and chemical characterization necessary when comparing populations.

Molecular analyses revealed genetically, morphologically and ecologically distinct subpopulations among cells traditionally assigned to *A. oxaliferum* [2, 4, 10]. In Lake Stechlin samples, 177 distinct 16S rRNA gene amplicon variants were recovered from only 22 freshwater cells [17], complicating interpretation of this marker. Individual cells are highly polyploid and contain divergent chromosome populations [8, 17, 18]. Nucleotide- and amino-acid-identity comparisons can place individual cells from the same sample [8], as well as sequences from the same cells [18] outside conventional single-species boundaries. Intracellular diversity therefore challenges both phylogenetic reconstruction and bacterial species delimitation.

Currently available genomes come from one freshwater, and two saline environments Mineral Springs, Florida (ca. 18 PSU), and the Sippewissett salt-marsh (ca. 35 PSU) populations [5, 17–19], but sampling remains uneven. The earlier comparative dataset contained only four partial saline single-cell genomes, whereas freshwater information included multiple single-cell and metagenomic datasets from Lake Stechlin [8, 18]. Consequently, apparent habitat associations may also reflect geography, lineage history and differences in genome recovery. The lack of sufficient data from saline environments led to a data-mining-based broad comparison of metabolic potential of the genus across different environments, suggesting a globally identical metabolic potential of the genus [8]. Additional marine single-cell genomes are needed to test if this is indeed the case, and determine whether *Achromatium* cells exhibit intracellular genomic diversity across different environments. These data would also allow us to assess whether such diversity coexists with differentiation among populations in both sequence relationships and functional potential.

Caribbean marine mangrove sediments provide an ideal sulfide-rich habitat for this comparison. Organic-matter degradation and sulfate reduction support thiotrophic symbioses [20–24], and diverse large sulfur-oxidizing bacteria [25], including *Beggiatoa* spp. [26], *Thiovulum* spp. [27–29], *Halochromatium* sp. [30] and the largest known species *Ca.* Thiomargarita magnifica [31]. Here, we identified a new population of *Achromatium* from these sediments. We combine light and electron microscopy, chemical microanalysis, and metagenomics and single-cell genomics of Guadeloupe mangrove cells. We compare these with freshwater and other saline references and ask whether population boundaries persist in the *Achromatium* genus despite the extensive diversity observed in some individual cells.

## Materials and Methods

### Sampling, microscopy and chemical analyses

Surface sediment beneath *Beggiatoaceae* mats was collected with a 50 mL syringe at approximately 1 m water depth in Manche-à-Eau lagoon, Guadeloupe (16°16′36″ N, 61°33′24″ W). Sediment was sieved through 90 µm mesh. Light microscopy, SEM–EDX and TEM characterized morphology, inclusion distributions and ultrastructure. SEM preparation used acetone dehydration, critical-point drying and gold coating [30], following fixation in 4% paraformaldehyde; imaging used an FEI Quanta 250 at 20 kV. EDX mapping followed the protocol described previously [30], using separately prepared cells fixed for 15 min at room temperature in slightly hypo-osmotic 2.5% glutaraldehyde and an Oxford M-max 50 mm² detector.

For TEM, we used conventional fixation, dehydration, resin-embedding and ultrathin sectioning [26], with the following conditions: 4% paraformaldehyde fixation for 2 h at 4 °C in filtered seawater; embedding in 1% low- melting-point agar; washing in 0.1 M sodium cacodylate, pH 6.8, at 900 mOsm; and 45 min postfixation in 1% osmium tetroxide and 1% ferrocyanide in the same buffer. Water-washed samples were ethanol-dehydrated, infiltrated with LR White under vacuum and polymerized for 48 h at 60 °C. Sections of 60 nm were examined with a Hitachi H7700 at 80 kV.

Raman spectra were acquired from visible inclusions in 4% paraformaldehyde-fixed cells maintained hydrated in filtered seawater, using a mIRage optical photothermal infrared (O-PTIR) microscope equipped with a Raman module (Photothermal Spectroscopy Corp., Santa Barbara, CA) with a 532 nm excitation laser through a 40x dry objective (NA = 0.78). Spectra were acquired over the range 180 to 3,300 cm-1 with three accumulations of 10 seconds each at 3.5% laser power (5 mW). Spectral preprocessing was performed in R (v4.4.2) using the Alkahest package [32]. Spectra were smoothed using the Savitzky-Golay algorithm [33], followed by background subtraction and baseline correction using the rolling ball method [34]. Spectra were compared with elemental sulfur and calcium-bearing references (RRUFF data base ID R040135 and R040170, respectively).

### Sequencing and assembly

Filtrate from the 90 µm mesh sieve with many *Achromatium* cells was used for cell purification on a sucrose gradient (750 g.L^-1^ sucrose; 0.05 M Tris-HCl; pH 7.4, 4°C) at 700 g for 3 min at 4 °C. The fraction containing large white cells, morphologically identified as *Achromatium* was collected. Individually micropipetted cells were further washed in three successive baths of 0.1 µm-filtered, UV-treated seawater and transferred into a 384-well plate. Three types of *Achromatium* samples were prepared for sequencing, i) single cells of the small morphotype (n=4), ii) single cells of the large morphotype (n=7), and iii) mixes of 5 to 10 cells of both morphotypes (n=4). The plate was shipped frozen to the Joint Genome Institute for sequencing. In short, water was removed, cells were lysed with lysozyme and whole-genome amplified by primary template-directed amplification (ResolveDNA Kit v2.0, BioSkryb Genomics). Libraries were prepared from 1 ng of DNA with the Nextera XT kit (Illumina) and sequenced on an Illumina NovaSeq X Plus (2 × 150 bp). Reads were normalized with BBTools v39.80 and assembled with SPAdes v4.2.0 (--sc). Contigs shorter than 2 kb or with insufficient read support were removed (see Supplementary Methods for details). The fifteen assemblies (4 small cells, 7 large cells, and 4 mixed-size) were analysed.

In addition, a large population containing thousands of *Achromatium* cells was used for DNA extraction and processed for metagenomics (see Heidig et al. [12] for details).

### Genome screening and marker-gene multiplicity

To decontaminate the assemblies from PTA-amplified cells, contig taxonomy was assigned with Kraken 2 (v2.17.1) using default parameters [35] against GTDB R226, and only *Achromatium* assignments were retained resulting in a “cleaned” single-cell assembly. Contig classification statistics are provided in Table S1. The metagenomic data was obtained earlier and used in Heidig et al. [12] and is referred to here as metagenomic bin1 and bin3. Limited reference representation may exclude divergent *Achromatium* contigs. Completeness, contamination and strain heterogeneity were assessed with CheckM1 1.2.5 [36] and CheckM2 1.1.0 [37]; ANI was calculated with fastANI 1.34 [38]. Per single-cell assembly, we recorded distinct markers recovered, multicopy-marker counts and maximum copies per marker. Neither measure estimates chromosomes completeness or independently validated loci. The bac120 marker genes form part of the functional gene repertoire but are analyzed separately because they are generally present once per haploid bacterial genome. Their expected single-copy status provides a reference for assessing gene multiplicity, while their conservation supports phylogenetic comparisons.

16S rRNA genes annotated by IMG in the complete single-cell assemblies, before Kraken 2 screening, were classified by megablast against NCBI core_nt. A single *Achromatium* 16S rRNA gene was found in these assemblies (type A) and served as the reference for the read analyses. To find additional 16S copies that the assemblies might have collapsed or missed, reads sharing a 20-mer with type A were extracted from each cell and mapped with BBMap 40.05 to all 16S rRNA genes found in the assemblies. In Small_H13, most of these reads were less than 97% identical to type A. Reads related to these divergent fragments were recruited from all Small_H13 reads by two rounds of k-mer baiting with BBDuk (k = 25) and assembled with SPAdes 4.3.0 (-- sc), yielding a second sequence (type B). After deduplication, reads assigned to one type at ≥97% identity, excluding ties, were counted as type-informative. Variable positions were called from reads ≥97% identical to the main type when the minor allele reached 2% and five reads (base quality ≥20). Reads not explained by the cell’s assembly (≥98% identity over ≥95% of their length; minimap2 2.28) were reassembled with SPAdes 4.3.0 (--sc) and searched for 16S rRNA genes with barrnap 1.10.5. For validation purposes, all 16S rRNA genes were additionally classified using SINA [39] with the SILVA release 144 database [40]. For phylogenetic analysis, sequences from this study were aligned with selected reference sequences from SILVA release 144 and Karst et al. [41], including nearest neighbours and sequences from Lake Stechlin metagenomes [17], using SINA 1.7.2. An approximately maximum-likelihood tree was inferred using FastTree 2 under the GTR model with Shimodaira–Hasegawa-like local support values [42]. The tree was rooted with *Allochromatium vinosum*, visualized using FigTree 1.4 [43], and its labels and colours were edited in PowerPoint (Fig. S8).

### Functional annotation and comparisons

Annotation used eggNOG-mapper 3.0.0 [44] against eggNOG v7 [45]. Five presence/absence matrices for 63 assemblies comprised 4,375 KEGG Orthology identifiers (KOs), 1,615 Enzyme Commission (EC) identifiers, 64 CAZy families, 3,710 PFAM families and 1,804 bacterial eggNOG groups. Copy number information per assembly was not used for the functional interpretation of the data. The primary comparison used 11 Guadeloupe single cells from this study and six Stechlin single cells previously sequenced [17]. Split sister-cell and combined cell-part assemblies [18], metagenomic bins [12, 17] and Guadeloupe mixed-cell assemblies provided supporting evidence. For downstream analyses we required sets to contain ≥800 distinct KOs. Subsequently from the split-sister-cells and combined-cell-part sets only nine and ten assemblies, respectively. WMS1–3 [5] and *Candidatus A. palustre* [19] were examined individually rather than pooled with Guadeloupe; the ordination analyses included 56 assemblies.

Jaccard dissimilarities were used for PCoA and one-factor PERMANOVA [46]. Two-sided Fisher exact tests compared feature-detection frequencies. Benjamini–Hochberg correction [47] was applied within annotation types for feature tests and across the five types for global tests (q < 0.05). Descriptive candidates required detection in ≥80% of one primary population and ≤20% of the other. Morphotype comparisons used seven large and four small cells; selected contrasts were exploratory. Population, site and study origin coincided, precluding experimental attribution to salinity. Analyses used Python 3.12.14, NumPy 2.3.5, pandas 2.2.3, SciPy 1.17.0 and Matplotlib 3.10.8.

### Phylogenetic inference and marker gene copy selection

The dataset comprised 63 *Achromatium* assemblies and *Allochromatium vinosum* as an outgroup. The bac120 markers were identified in both the new and existing datasets using GTDB-Tk 2.7.2 [48] with GTDB R232 [49]. GTDB-Tk’s trimmed bac120 tree retained 53 *Achromatium* assemblies; ten partial-cell assemblies had <75% of markers. For copy-selection analyses we required ≥90 bac120 families and ≥500 eggNOG-named proteins. All marker copies were retained. Following median-based length filtering, families present in ≥90% of assemblies were aligned with MAFFT 7.526 L-INS-i [50]. Families with >1% unmeasurable pairwise distances (insufficient overlap or Kimura p > 0.854) were removed. One hundred independent concatenations selected one copy uniformly per marker and assembly from fixed alignments, with gaps for missing markers.

Maximum-likelihood trees were inferred using IQ-TREE 3.1.2 under LG+G4 with --keep-ident [51]; for the displayed supports 1,000 UFBoot and 1,000 SH-aLRT replicates were used [52, 53]. All copies of each individual marker were used to infer the corresponding single-marker trees. Additional trees were inferred from random-copy concatenations, GTDB-Tk’s trimmed alignment, and GTDB-Tk-selected copies realigned using the same procedure as for the random-copy dataset. The trimmed tree was outgroup-rooted, individual-marker trees unrooted and replicate trees rooted on the freshwater clade. Normalized Robinson–Foulds distances and sister-group changes were measured relative to replicate 0. Marker-tree environmental separation was the minimum copies requiring repositioning to separate freshwater and saline samples by one branch. Trees were parsed with ete4 4.4.0 and drawn with Matplotlib 3.11.1 in Python 3.14.7.

## Results

### Cell morphology and inclusion chemistry

Structural characterization of the purified *Achromatium* cells showed co-occurring small and large morphotypes with mean sizes of 14 ± 2 µm and 22 ± 3 µm, respectively. Spherical to ellipsoidal cells had rounded surface reliefs corresponding to underlying inclusions (Fig. 1A–D). TEM distinguished larger and smaller inclusion profiles, averaging 804 ± 118 nm and 155 ± 32 nm, with smaller inclusions distributed between larger ones (Fig. 2A, B; asterisks and arrows, respectively). The larger inclusions occupied much of the cell volume and showed low contrast without visible crystalline structures. Surrounding membrane-delimited regions contained approximately 22 nm electron-dense structures compatible in size with ribosomes. The cytoplasm was highly compartmentalized, with an extensive network of intracytoplasmic membranes delimiting numerous internal compartments with an average size of 617 ± 190 nm (n = 10; Fig. 2C). At the cell periphery, two distinct membrane profiles were visible, consistent with a diderm cell envelope (Fig. 2D).

**Figure 1.**
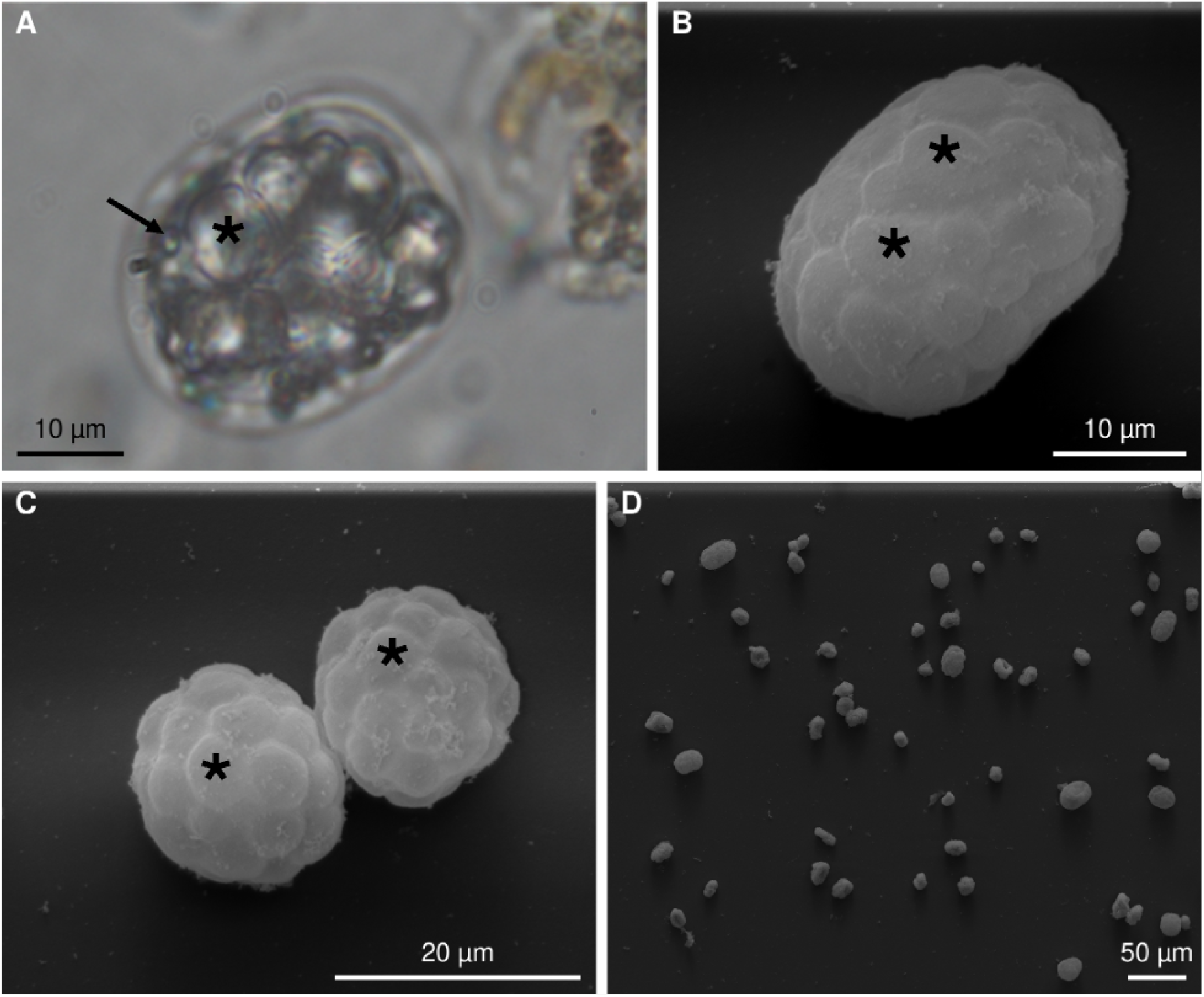
Structural characterization of *Achromatium* cells from Caribbean mangrove sediments. (A) Light microscopy image of an *Achromatium* cell showing numerous intracellular inclusions. (B) SEM image of the large morphotype. (C) SEM image of the small morphotype. (D) SEM image showing the coexistence of multiple *Achromatium* cells within the sampled population. Black asterisks indicate large inclusions (A) or their corresponding surface reliefs (B, C); the black arrow in A indicates a smaller inclusion.

**Figure 2.**
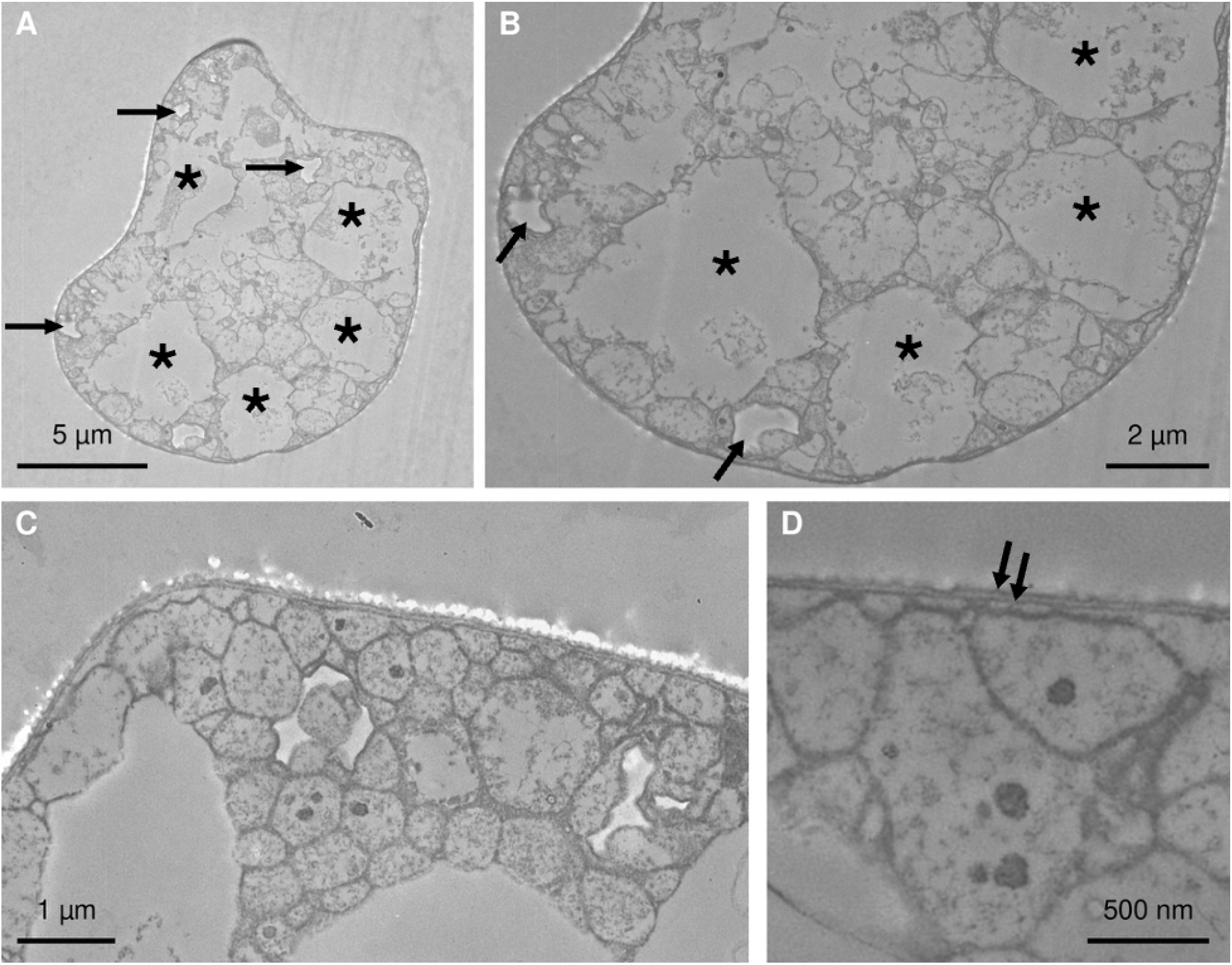
Intracellular ultrastructure and cell envelope of *Achromatium* cells observed by transmission electron microscopy. (A) Overview of a cell section containing large low-contrast inclusion profiles (asterisks) and smaller inclusion profiles (arrows). (B) Higher-magnification view of the cell shown in A, illustrating the distribution of large and small inclusion profiles. (C) Peripheral cytoplasm showing extensive compartmentalization by intracytoplasmic membranes and numerous electron-dense structures. (D) Higher-magnification view of the cell wall; paired arrows indicate the two membrane profiles consistent with a diderm organization.

SEM–EDX detected spatially distinct calcium-rich and sulfur-rich regions: calcium was broadly associated with larger inclusions, whereas sulfur formed smaller spots (Fig. 3). Raman bands at 222 and 476 cm⁻¹ supported elemental sulfur, but spectra did not identify the calcium-rich inclusions as the tested crystalline calcium phases (Fig. S1). Their mineralogical identity therefore remained unresolved. Together, the observed cell morphology, ultrastructure and calcium- and sulfur-rich inclusions were consistent with assignment to the genus *Achromatium*.

**Figure 3.**
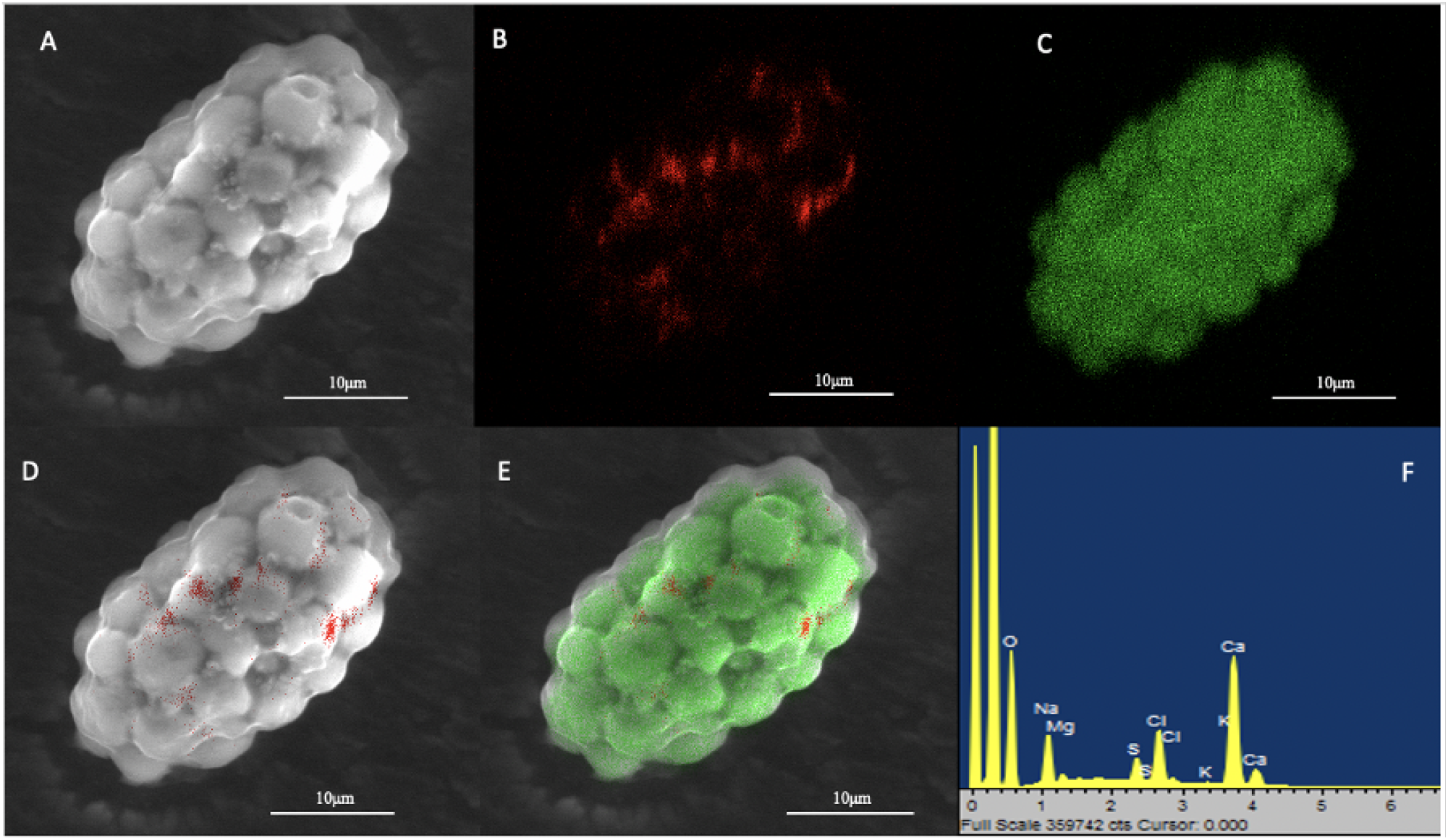
Sulfur and calcium distribution in an *Achromatium* cell analysed by SEM–EDX. (A) Scanning electron micrograph of the analysed cell. (B–C) Elemental maps showing the spatial distribution of sulfur (B, red) and calcium (C, green). (D–E) Mixmaps obtained by superimposing the sulfur map (D) or both sulfur and calcium maps (E) onto the SEM image. (F) Energy-dispersive X-ray spectrum of the analysed cell area, showing the detected elements, including sulfur (S) and calcium (Ca).

### Assembly properties and nucleotide identity

The 11 screened single-cell assemblies contained 338–889 contigs and 3.9–6.7 Mb. CheckM1 and CheckM2 completeness estimates ranged from 77.2–100.0% and 90.8–100.0%, respectively. Nine assemblies had low contamination estimates; Large_I06 and Small_H13 had substantially higher contamination estimates from both tools (Table 1) despite consisting exclusively of contigs identified as *Achromatium*.

**Table 1.** Assembly statistics and CheckM quality estimates for the 11 marine mangrove *Achromatium* single-cell assemblies after contig screening. Comp., completeness; Contam., contamination; strain het., strain heterogeneity.

| Single-cell ID | Contigs | Length (Mb) | CheckM1 Comp. (%) | CheckM1 Contam. (%) | CheckM1 Strain het. (%) | CheckM2 Comp. (%) | CheckM2 Contam. (%) |
|---|---|---|---|---|---|---|---|
| Large_M02 | 377 | 4.03 | 84.96 | 1.02 | 0.00 | 96.06 | 0.29 |
| Large_L04 | 701 | 4.08 | 86.24 | 2.21 | 8.33 | 96.51 | 1.95 |
| Large_M04 | 889 | 4.22 | 85.55 | 1.52 | 16.67 | 96.29 | 0.49 |
| Large_H05 | 363 | 4.01 | 84.22 | 0.79 | 0.00 | 95.27 | 0.12 |
| Large_J05 | 346 | 3.99 | 84.98 | 0.75 | 0.00 | 94.34 | 0.90 |
| Large_I06 | 487 | 5.18 | 86.75 | 29.86 | 5.81 | 98.48 | 28.75 |
| Large_N11 | 587 | 3.91 | 77.17 | 2.18 | 11.76 | 90.84 | 0.59 |
| Small_H13 | 671 | 6.70 | 100.00 | 100.00 | 31.58 | 100.00 | 72.58 |
| Small_J13 | 370 | 4.14 | 83.99 | 1.16 | 0.00 | 94.05 | 0.42 |
| Small_J14 | 365 | 4.03 | 84.17 | 1.18 | 16.67 | 97.01 | 1.34 |
| Small_H15 | 338 | 4.10 | 85.44 | 0.80 | 0.00 | 96.03 | 0.15 |

All retained contigs were assigned to *Achromatium* by Kraken 2 against GTDB R226. Estimates describe the screened assemblies and do not independently validate chromosome-copy number.

ANI among Guadeloupe single-cell assemblies ranged from 96.2–99.9%. Across all 17 Guadeloupe assemblies, including mixed-cell assemblies and bins, ANI ranged from 77.3–99.9% over 136 comparisons, with median 98.2%. The lowest identities primarily involved metagenomic Bin3; Small_H13 also differed from the remaining assemblies. Within Stechlin, 815 available comparisons ranged from 73.4–99.9% (median 81.5%). The 712 Guadeloupe–Stechlin comparisons ranged from 73.4–82.6% (median 74.2%; Fig. 4).

**Figure 4.**
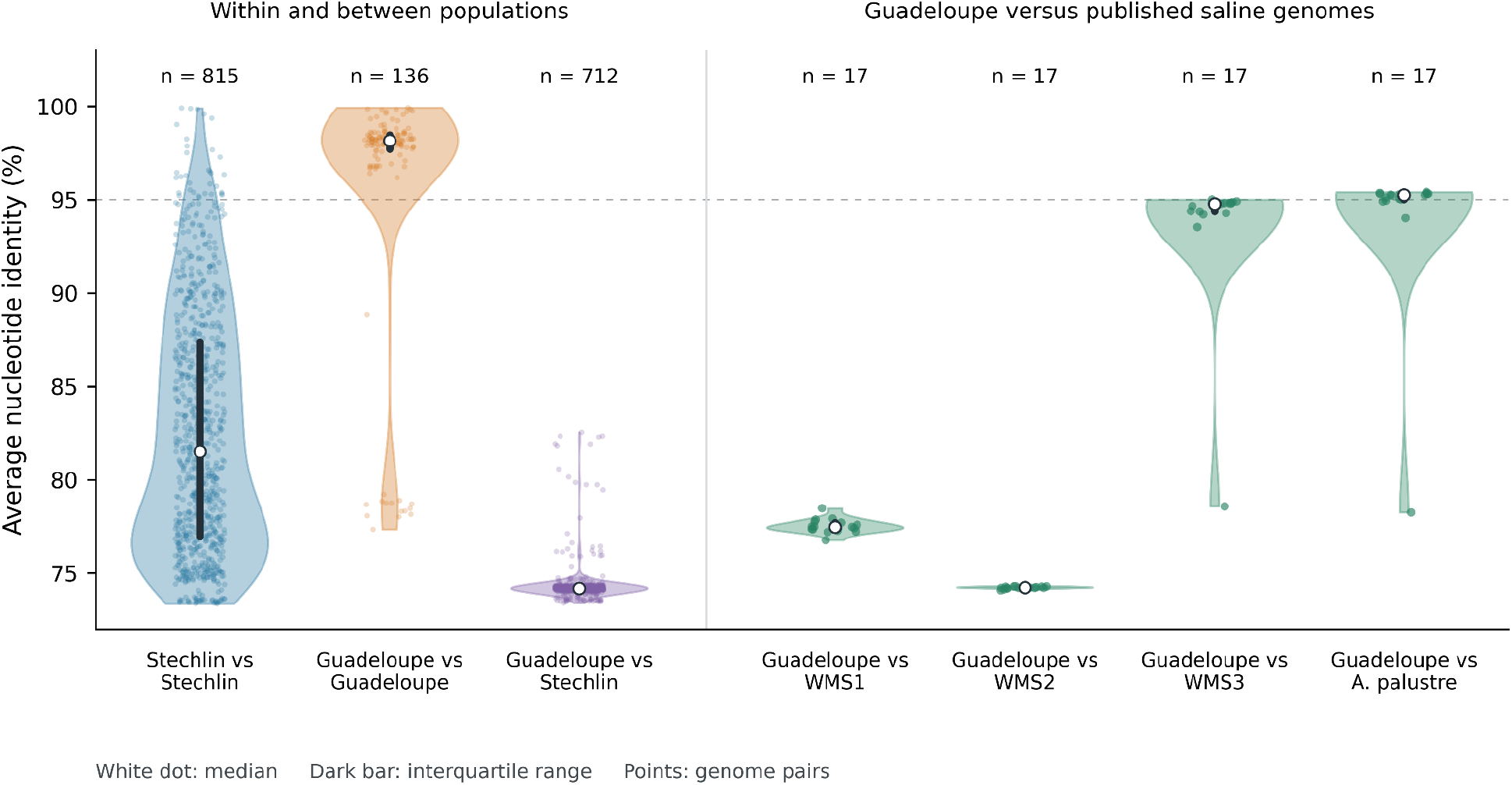
Average nucleotide identity among *Achromatium* genome assemblies. Seven violin plots summarize Stechlin– Stechlin, Guadeloupe–Guadeloupe, Guadeloupe–Stechlin, and separate Guadeloupe comparisons with WMS1, WMS2, WMS3 and *Candidatus* Achromatium palustre. The numbers above the violins indicate available pairwise ANI values. Violin widths show Gaussian kernel density estimates with Scott bandwidth, scaled separately to equal maximum widths and bounded by the observed range. Points show all available genome pairs with horizontal jitter; white dots indicate medians and dark bars the interquartile ranges. The dashed line marks 95% ANI as a visual reference.

Each published saline reference yielded 17 comparisons with Guadeloupe. ANI ranges and medians were 76.8–78.5% and 77.5% for WMS1; 74.1–74.3% and 74.2% for WMS2; 78.6–95.0% and 94.8% for WMS3; and 78.3–95.4% and 95.3% for *Candidatus* A. palustre. The lowest values against WMS3 and *Candidatus* A. palustre involved metagenomic Bin3 (Fig. 4).

### Functional repertoires of Achromatium populations

The Guadeloupe cells contained 1,323–1,488 distinct KOs per assembly, compared with 1,398–1,820 in the six single cells from Lake Stechlin [17]. Their combined recovered repertoires contained 1,777 and 2,308 KOs, respectively, of which 1,396 were shared. Consistent differences between Guadeloupe and Stechlin populations were also detected using EC identifiers, CAZy families, PFAM families and bacterial-level eggNOG orthologous groups (PERMANOVA, all q < 0.001; Fig. S4). In contrast, none of the five annotation types showed a significant difference between the small and large Guadeloupe morphotypes after multiple-testing correction. KO-based PCoA separated the Guadeloupe assemblies from both the Stechlin single cells and the split-sister-cell-series assemblies (Fig. 5A). Distinct KO recovery increased with the number of predicted proteins recovered (Fig. 5B). In the primary comparison, median Jaccard dissimilarity was 0.43 between the Guadeloupe and Stechlin populations, compared to 0.14 within Guadeloupe and 0.32 within Stechlin. KO profiles were associated with population in the exact permutation test (pseudo-F = 24.72, R² = 0.62, p = 8.08 × 10⁻⁵; q = 8.08 × 10⁻⁵ across the five annotation-type tests). KO dispersion differed between populations as well (permutation p = 0.037), with greater within-population dissimilarity in Stechlin; the PERMANOVA association therefore did not represent a test of centroid separation alone. At the individual-feature level, 524 KO annotations were associated with a specific population after false-discovery-rate correction. The descriptive prevalence screen identified 185 Guadeloupe-associated and 229 Stechlin-associated KOs.

**Figure 5.**
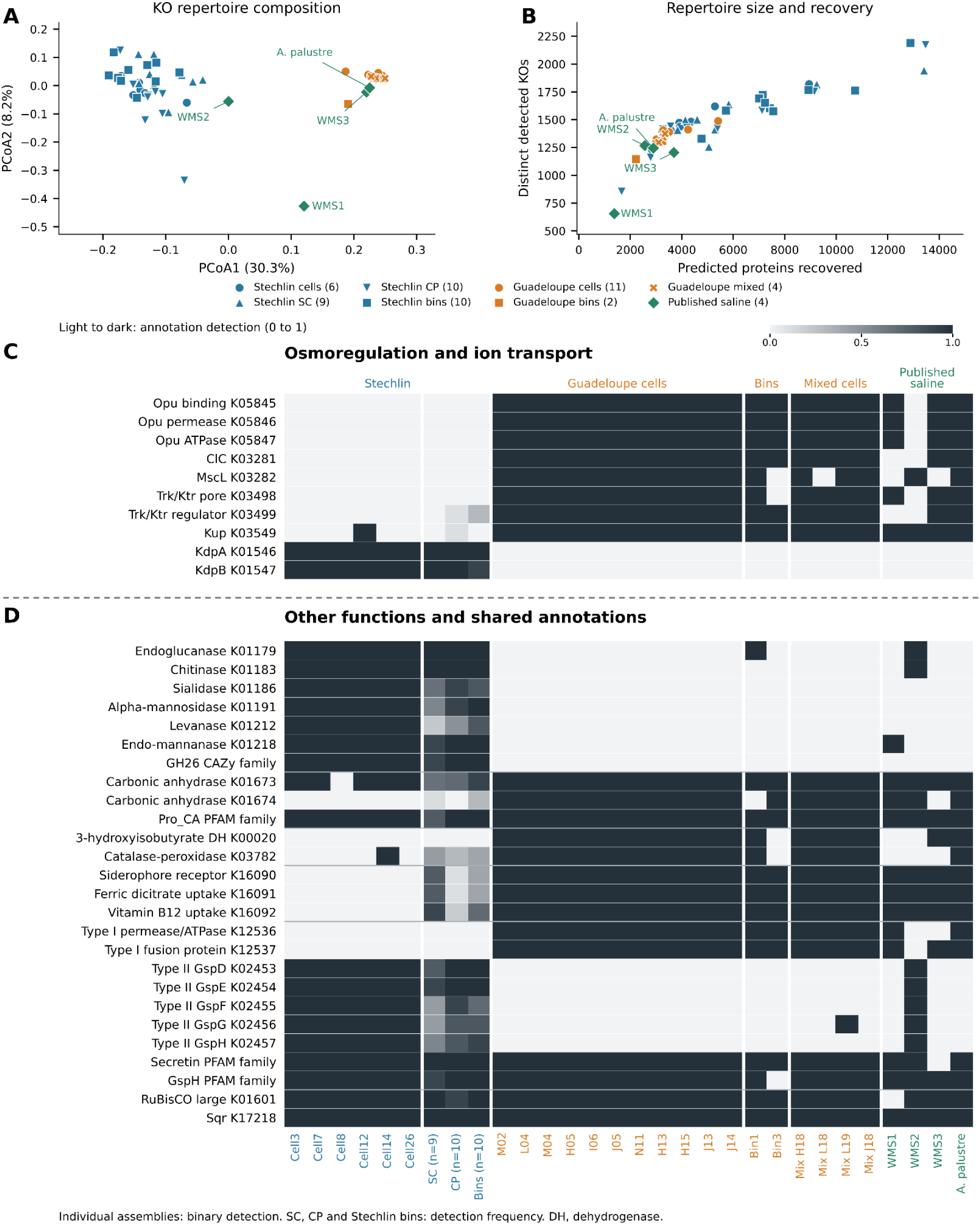
Functional comparison of Guadeloupe, Lake Stechlin and published saline-environment *Achromatium* genomes. (A) PCoA of binary KO profiles using Jaccard dissimilarity, including all 56 assemblies represented in panels C and D. Each assembly was included individually; group sizes and symbols are given in the legend. SC denotes split-cell assemblies; CP denotes merged data from all sequenced parts of an individual cell (SC and CP taken from [18]). Colours identify Stechlin (blue), Guadeloupe (orange) and published saline genomes (green). Axis percentages refer to positive PCoA inertia. (B) Distinct KO recovery versus predicted protein recovery for the same assemblies. (C) Selected osmoregulatory and ion-transport annotations. (D) Selected carbohydrate-processing, carbonic anhydrase, metabolic, nutrient-acquisition and secretion annotations, with shared metabolic markers. The dashed line separates panels C and D. Light and dark cells denote non-detection and detection in individual assemblies; intermediate shading in the SC, CP and Stechlin-bin summary columns denotes detection frequency across 9, 10 and 10 assemblies, respectively. Pro_CA is shared by every primary cell; K01673 is undetected in Cell8. Secretion rows show selected components and families rather than complete systems. Primary population tests compare 11 Guadeloupe with six Stechlin single cells. Shading represents detection rather than copy number; non-detection does not establish biological absence.

The published saline-environment genomes placed the Guadeloupe results in a broader comparative context (Fig. 5A, B). *Candidatus* A. palustre and WMS3 were more similar to Guadeloupe than to the primary Stechlin set: their median KO-based Jaccard dissimilarities to Guadeloupe were 0.21 and 0.25, respectively, compared with 0.45 and 0.48 to Stechlin. WMS2 showed similar dissimilarities to the two populations (0.457 and 0.467), whereas WMS1 was more dissimilar to both (0.62 and 0.68). From WMS1 only 655 distinct KOs were recovered, compared with 1,267 from WMS2, 1,205 from WMS3 and 1,244 from *Candidatus* A. palustre; its more distant profile therefore coincided with substantially lower annotation recovery.

### Transport and metabolic annotations

The clearest contrasts involved solute and ion transport (Fig. 5C). All 11 Guadeloupe cells carried the three Opu osmoprotectant-transport genes (K05845–K05847), the ClC-family (K03281), the mechanosensitive channel (K03282) and the Trk/Ktr K03498 and K03499. None was detected in the six Stechlin single cells (q = 0.001 in the KO screen). Kup K03549 was detected in all Guadeloupe cells and one Stechlin cell (q = 0.007). Conversely, KdpA and KdpB, (K01546 and K01547), occurred in all six Stechlin single cells and no Guadeloupe cells (each q = 0.001). Opu, ClC, MscL and KdpA/B contrasts retained the same direction in the recovery-filtered SC and CP sets. The Guadeloupe metagenomic bins and mixed-cell assemblies broadly reproduced the single-cell transporter profile, while published saline genomes showed similar but variable patterns (Fig. 5C). Similar, but not identical, transporter profiles were observed in the published saline genomes (Fig. 5C)

Functional contrasts extended beyond osmoregulation (Fig. 5D). Six carbohydrate-processing KOs, encoding endoglucanase (K01179), chitinase (K01183), sialidase (K01186), alpha-mannosidase (K01191), levanase (K01212) and endo-beta-mannanase (K01218), were each detected in all six Stechlin single cells and no Guadeloupe single cells (each q = 0.00110). Endoglucanase and chitinase were also detected in all nine supporting SC assemblies, all ten CP assemblies and all ten Stechlin metagenomic bins. CAZy GH26, GH5 and CBM11 occurred in all six Stechlin single cells and none of the Guadeloupe cells, bins or mixed-cell assemblies. The GH26 contrast was also recovered in the PFAM-family and EC 3.2.1.78 profiles. These differences concerned particular annotation assignments: for example, EC 3.2.1.14 and EC 3.2.1.18 were each recovered from one Guadeloupe cell despite non-detection of K01183 and K01186, respectively.

Carbonic anhydrase profiles distinguished a shared family-level signal from an additional Guadeloupe-associated ortholog (Fig. 5D). K01673 was detected in all 11 Guadeloupe cells and five of the six Stechlin single cells. The related Pro_CA PFAM family was detected in every primary cell, including Stechlin Cell8, which lacked the K01673 assignment. In contrast, K01674 and the Carb_anhydrase family were detected in all Guadeloupe cells and none of the six Stechlin single cells (K01674 q = 0.001). K01674 was also detected in some of the Stechlin supporting assemblies, so its distribution was not exclusive to Guadeloupe across the complete dataset. K01673 and Pro_CA were recovered in both Guadeloupe bins, all four mixed-cell assemblies and all four published saline genomes. K01674 occurred in all four other saline assemblies other than Bin1 and WMS3.

Other Guadeloupe-associated KOs included 3-hydroxyisobutyrate dehydrogenase (K00020), associated with branched-chain amino acid degradation, and catalase-peroxidase (K03782). K00020 was detected in all 11 Guadeloupe cells, all four mixed-cell assemblies, Bin1, WMS3 and *Candidatus* A. palustre, but in none of the primary Stechlin single cells, recovery-filtered SC or CP assemblies or Stechlin bins (q = 0.001 in the primary comparison). K03782 occurred in all Guadeloupe single-cell and mixed-cell assemblies, Bin1 and *Candidatus* A. palustre, compared with one of six original Stechlin single cells (Cell14; q = 0.007) and a subset of the supporting Stechlin assemblies. Its distribution indicated higher prevalence in Guadeloupe rather than population exclusivity.

The catecholate siderophore receptor K16090, ferric dicitrate transporter (K16091) and vitamin B12 transporter (K16092) were each detected in every Guadeloupe from both Guadeloupe bins, all four mixed-cell assemblies and all four published saline genomes. However, in Stechlin K16090 and K16091 occurred in seven of nine supporting Stechlin SC assemblies, and K16092 in eight of nine but not in the six Stechlin single cells (each q = 0.001). Secretion-associated annotations showed contrasting distributions (Fig. 5D). The type I secretion permease/ATPase (K12536) and membrane fusion protein K12537 occurred in all Guadeloupe cells and none of the six Stechlin single cells (each q = 0.001). Both were also detected in the two Guadeloupe bins and all four mixed-cell assemblies, but in none of the recovery-filtered SC or CP assemblies or Stechlin bins. Both occurred in WMS1 and *Candidatus* A. palustre; WMS3 carried K12537 alone, and neither was detected in WMS2. Conversely, 11 type II general secretion pathway annotations (K02451, K02453–K02458 and K02460–K02463) were each recovered in all six Stechlin single cells and no Guadeloupe single cells (each q = 0.00110). None was detected in either Guadeloupe bin; mixed sample L19 carried GspG K02456 alone among these 11 annotations. WMS2 carried ten of the 11, whereas none was detected in WMS1, WMS3 or *Candidatus* A. palustre. Nevertheless, the Secretin and GspH PFAM families were shared by every primary cell. The KO patterns thus described differences in assigned secretion components and did not establish the absence of all related machinery.

These contrasts occurred alongside shared metabolic markers. RuBisCO large and small subunit annotations occurred in every primary cell, both Guadeloupe metagenomic bins and all four mixed-cell assemblies; phosphoribulokinase occurred in ten Guadeloupe cells and all six Stechlin single cells. Sqr and several Sox-associated annotations were widely shared. Sqr was detected in all four published saline genomes, while RuBisCO large and small subunits were detected in WMS2, WMS3 and *Candidatus* A. palustre but not WMS1. The Guadeloupe bins also retained some differences from the single-cell pattern: Bin1 carried endoglucanase (K01179), whereas the other five Stechlin-associated carbohydrate KOs were undetected in both bins and all four mixed-cell assemblies. Thus, the supporting assemblies reproduced many recurring signals without showing an identical annotation inventory. These marker detections were not evaluated as complete pathway reconstructions.

### Functional overlap between morphotypes of Guadeloupe isolates

The large and small Guadeloupe cells had broadly overlapping functional repertoires (Fig. S2). Large cells contained 1,323–1,411 distinct KOs and small cells 1,347–1,488; without Small_H13, the small-cell range was 1,347–1,405. KO profiles showed a modest nominal association with morphotype (pseudo-F = 1.80, R² = 0.17, p = 0.042), but this did not meet the significance threshold after correction across annotation types (q = 0.071). None of the five global annotation-type comparisons, or any individual feature-level contrast, remained significant at q < 0.05 (Figs. S2 and S5).

Three exploratory KO contrasts were highlighted (Fig. S2C): fructokinase K00847 was detected in six of seven large cells and no small cells, whereas ThiH K03150 and the S-adenosylmethionine uptake-transporter annotation K15270 were each detected in three of four small cells and no large cells. All three had feature-level q = 1.00 and therefore were not statistically supported morphotype markers. No KO occurred in every cell of one morphotype and was absent from every cell of the other. Small_H13 was the most distinct small-cell profile and contained 95 KOs undetected in the other ten cells. Nevertheless, removing it did not eliminate the nominal KO-level association. The Opu, ClC, MscL and Trk/Ktr annotations characteristic of the Guadeloupe comparison were recovered in both morphotypes.

### Intracellular genomic diversity revealed by conserved marker genes and 16S rRNA types

Between 110 and 116 of the 120 GTDB bac120 marker families were recovered per single-cell assembly (Table 2). Nine assemblies contained two or three multicopy markers, each represented by two copies. Large_I06 contained 33 multicopy markers with up to three copies per marker, whereas Small_H13 contained 107 multicopy markers among 114 recovered families, with up to six copies per marker. Elevated marker multiplicity therefore occurred in one cell of each morphotype (Table 2; Fig. S6). Within-assembly comparisons revealed sequence divergence between copies of GTDB bac120 markers, including in Large_I06 and Small_H13 (Fig. S7). Analysis of the 16S rRNA gene in the assemblies showed the same split. Ten cells each contained a single *Achromatium* 16S rRNA gene (16S type A, 99.5% identical to *Ca.* A. palustre), identical among cells apart from one substitution in Large_J05, while the Small_H13 assembly contained none. In the reads, nine cells carried only this sequence type, apart from two or three variable positions in Large_J05 and Small_J14. Large_I06 and Small_H13 also contained a second type (type B), 93.8% identical to the first, which made up 3.5% and 89% of type-informative reads, respectively. Reassembling the reads that the assemblies did not explain recovered the complete second type from Small_H13 and no other *Achromatium* 16S rRNA sequence. The expanded 16S rRNA gene phylogeny did not resolve a complete separation between freshwater and saline *Achromatium* sequences (Fig. S8).

**Table 2.** Recovery and multiplicity of GTDB bac120 markers in marine mangrove *Achromatium* single-cell assemblies. Multicopy markers are families represented by more than one recovered copy; maximum marker copies is the highest count for one marker in an assembly. These counts do not estimate complete chromosomes.

| **Single-cell ID** | **Morphotype** | **GTDB markers recovered** | **Multicopy GTDB markers** | **Maximum marker copies** |
|---|---|---|---|---|
| Large_M02 | Large | 116/120 | 3 | 2 |
| Large_H05 | Large | 114/120 | 2 | 2 |
| Large_I06 | Large | 114/120 | 33 | 3 |
| Large_J05 | Large | 114/120 | 3 | 2 |
| Large_L04 | Large | 114/120 | 2 | 2 |
| Large_M04 | Large | 113/120 | 2 | 2 |
| Large_N11 | Large | 110/120 | 3 | 2 |
| Small_H13 | Small | 114/120 | 107 | 6 |
| Small_H15 | Small | 114/120 | 2 | 2 |
| Small_J13 | Small | 113/120 | 2 | 2 |
| Small_J14 | Small | 114/120 | 3 | 2 |

The maximum-likelihood phylogeny reconstructed from GTDB-Tk’s trimmed bac120 alignment included 53 *Achromatium* assemblies and the *Allochromatium vinosum* outgroup. Most saline-associated assemblies, including the Guadeloupe *Achromatium* mangrove samples, clustered separately from freshwater references, whereas WMS2 did not join the principal saline-associated group (Fig. 6A). Individual marker-gene trees containing all recovered copies showed different relationships between the two sampling environments. Freshwater and saline copies were fully separated in arginyl-tRNA synthetase (TIGR00456) (Fig. 6B). 30S ribosomal protein S6 (TIGR00166) showed a broadly similar grouping, with a WMS2 copy outside the saline-associated group (Fig. 6C), whereas no single branch separated environments in ribosome-binding factor A (TIGR00082) (Fig. 6D). ClpX (TIGR00382) had the highest mean copy number among retained markers, at 2.1 average copies per assembly, and its tree also contained copies from different sampling environments within the same broader groupings (Fig. 6E). The expression of sspB variants, an adaptor protein to ClpX, were recently suggested to be temperature regulated [12].

**Figure 6.**
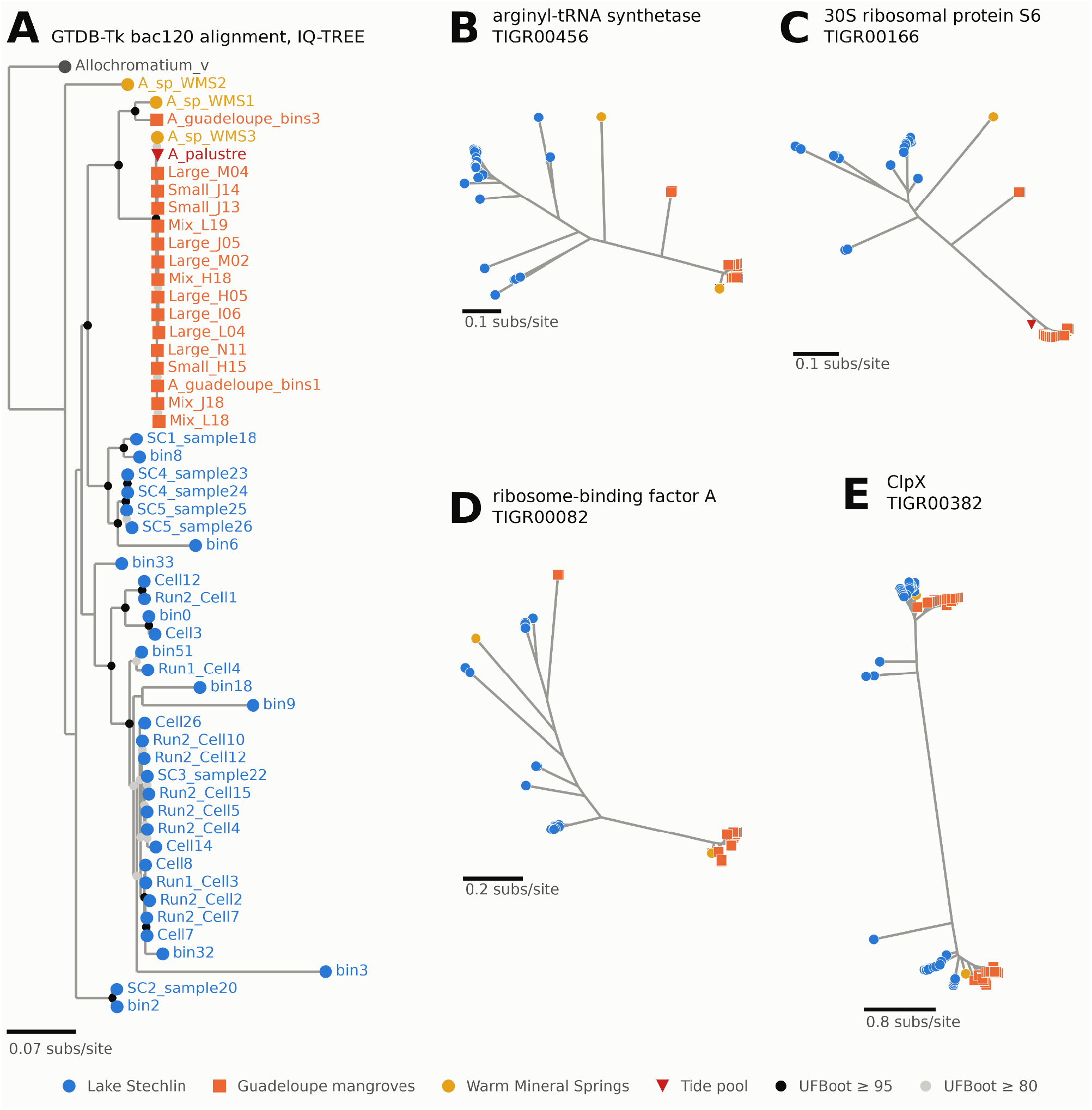
Phylogenies of *Achromatium* assemblies and copies of individual marker genes. Colours and tip symbols identify sampling origin: blue circles, Lake Stechlin; orange squares, Guadeloupe mangroves; yellow circles, Warm Mineral Springs; and red inverted triangles, the tide-pool reference. The *Allochromatium vinosum* outgroup is shown in grey. (A) Maximum-likelihood tree inferred from GTDB-Tk’s trimmed bac120 alignment for 53 *Achromatium* assemblies and rooted on *Allochromatium vinosum*. Ten *Achromatium* assemblies were excluded from this alignment. Grey and black node symbols indicate ultrafast bootstrap support of at least 80 and at least 95, respectively. (B–E) Unrooted maximum-likelihood trees of individual marker genes, with each recovered copy represented as a separate tip. (B) Arginyl-tRNA synthetase (TIGR00456), in which freshwater and saline copies are fully separated. (C) 30S ribosomal protein S6 (TIGR00166), illustrating separation of most copies by environment with a WMS2 copy outside the saline-associated grouping. (D) Ribosome-binding factor A (TIGR00082), for which no single branch separates copies from the two environments. (E) ClpX (TIGR00382), the most multicopy marker in the filtered dataset, with a mean of 2.1 copies per assembly. Very short branches in B–E were displayed at a minimum length to keep overlapping tips visible. Scale bars indicate substitutions per site.

The filtered copy-selection dataset contained 44 assemblies and 96 marker families represented by 4,883 copies. Selecting one copy per marker per assembly produced 100 alternative concatenated alignments of 40,668 amino-acid columns. Normalized RF distances between replicate 0 and the other 99 phylogenies ranged from 0.29 to 0.56, with a median of 0.44 (Fig. S3A–C, E). The tree reconstructed using GTDB-Tk’s copy selections, realigned and concatenated with the same procedure as the random-copy dataset, had an RF distance of 0.56 from replicate 0 (Fig. S3D–E).

The effect of copy selection differed markedly among assemblies. Seven changed sister group in at least 95% of alternative draws; six of these were from Guadeloupe. Six assemblies never changed sister group (Fig. S3F). The four cells belonging to the SC4 and SC5 sister-cell pairs formed a single clade in every draw, despite changes elsewhere in the trees (Fig. S3A–D). The illustrated tanglegrams retained the broad distinction between freshwater and saline-associated groups while showing changes in relationships within those groups.

## Discussion

The Guadeloupe *Achromatium* genomes broaden our understanding of intracellular genomic diversity in marine *Achromatium* and enable comparisons with freshwater cells from Lake Stechlin. These comparisons reveal population-associated differences in sequence and functional potential despite intracellular genomic diversity. Phylogenetic separation is clearer in multigene analyses than in individual gene trees. Together, these findings refine the view of a broadly conserved functional inventory and provide a basis for investigating how ecological specialization and speciation proceed alongside intracellular genomic diversity.

### Cell organization and mangrove habitat

The Guadeloupe population combines the large size, calcium-rich inclusions and localized sulfur characteristic of *Achromatium* [14, 15], extending cellular and genomic comparisons into Caribbean mangroves. Larger inclusions surrounded by smaller profiles resemble other populations, although the profiles measured here were smaller than the 3–6 µm carbonate bodies and approximately 1 µm sulfur globules reported for Stechlin lake [15]. Small inclusion profiles were regularly observed among approximately 50 examined cells. Unmeasured sediment pH and calcium concentrations prevent assessment of a geochemical explanation.

Membrane-delimited regions containing ribosome-sized structures are compatible with the complex organization described in freshwater cells, where periplasmic carbonate pockets partition the cytoplasm [15, 18]. However, our TEM images do not resolve the connectivity of these membrane-delimited regions or establish the precise localization of calcium-rich inclusions in the mangrove cells. Likewise, calcium enrichment, low TEM contrast and failure to match crystalline Raman references do not identify a mineral phase. The amorphous calcium carbonate demonstrated in freshwater *Achromatium* [16] is a plausible interpretation requiring direct confirmation.

Occurrence beneath Beggiatoaceae mats is consistent with access to opposing sulfide and oxygen gradients. Mangrove sediments can show oxygen penetration of approximately 1 mm [54], and active sulfate reduction has been documented in mangrove sediments covered by Beggiatoaceae mats in Guadeloupe [25]. Depth- and diel-dependent inclusion chemistry in salt-marsh *Achromatium* [14], and negative chemotaxis away from high oxygen, sulfide and nitrate concentrations in freshwater cells [13], suggest mechanisms for responding to variable redox conditions. Their operation in the mangrove population remains untested, as do alternative electron acceptors and changes in inclusion contents.

### Saline and freshwater *Achromatium* shared main metabolic pathways yet harbor population-specific functions

Annotation profiles of the different populations refine the proposed broadly conserved functional inventory of *Achromatium* which was based mostly on data mining from non-targeted metagenomic datasets [8]. Shared sulfur-oxidation and carbon-fixation markers coexist with differences in transport, carbohydrate processing, carbonic anhydrases and secretion-associated components. Guadeloupe Trk/Ktr and Kup annotations contrast with Stechlin KdpA/B, but do not indicate loss of potassium uptake. KdpFABC responds to potassium limitation even in the extreme halophile *Halobacterium salinarum* [55], whereas Trk-dependent uptake has been demonstrated in halophilic *Halomonas elongata* [56]. Opu, chloride-transporters and mechanosensitive-channels provide further candidates for ionic and osmotic regulation. Their recurrence in *Candidatus* A. palustre and WMS3 extends the comparison beyond Guadeloupe, but WMS2 shows that the full combination is not universal among saline genomes.

Carbohydrate-processing differences identify potential variation in organic-substrate use alongside shared autotrophic capabilities. Repeated Stechlin endoglucanase, chitinase, mannanase and GH5/GH26/CBM11 detections support targeted enzyme comparisons. Nevertheless, Guadeloupe EC detections and Bin1 endoglucanase prevent claims of population-exclusive carbohydrate utilization. Family assignments alone establish neither substrate specificity nor enzyme localization or rates. K00020 similarly identifies a candidate reaction in branched-chain amino-acid degradation, not a complete pathway or substrate preference. Catalase-peroxidase differences concern the prevalence of one peroxide-processing enzyme, not absence of freshwater oxidative-stress defences.

Both populations shared β-class carbonic anhydrases, while Guadeloupe consistently carried an additional α-class family. Because bacterial α-carbonic anhydrases can be periplasmic, this difference suggests potential variation in the cellular location of inorganic-carbon processing and pH regulation. Confirming the localization of these enzymes would help assess their possible relationship to the calcium-rich inclusions.

Differences in secretion-associated annotations also suggest variation in protein export and interactions with the surrounding sediment. However, the recovery of related components in both populations indicates that these differences concern particular components rather than entirely distinct secretion systems.

The metagenomic bins and mixed-cell assemblies broadly supported the functional patterns observed in single cells. However, these pooled datasets cannot establish that all detected genes occur together within an individual cell or define its metabolic repertoire.

### Population structure amid intracellular genomic diversity

The 16S rRNA gene did not show intracellular diversity for nine of the 11 single cells from Guadeloupe which carried a single 16S type, identical among cells, whereas individual Lake Stechlin cells were reported to carry several variants differing by up to 7% [17]. A second 16S type was restricted to Large_I06 and Small_H13. The recovery of divergent copies of conserved marker genes extends the evidence for intracellular genomic diversity to marine mangrove single cells, consistent with freshwater heterozygous polyploidy [8, 17]. While marker-copy counts cannot quantify complete chromosomes, the variation in marker multiplicity among the 11 single-cell assemblies indicates marked heterogeneity in the sequences recovered from individual cells. On the level of individual marker gene phylogenies, the separation of saline and freshwater environments is varied. The interspersion of freshwater and saline sequences in the 16S rRNA gene phylogeny (Fig. S8) further indicates that the environment-associated structure recovered with concatenated markers should not be interpreted as a universal separation across loci. Yet, the concatenated bac120 tree and illustrated replicates retained a broad association between sampling environment and phylogenetic grouping. Geographically restricted freshwater sampling prevents attribution of this structure to salinity alone. The case of WMS2 illustrates that a saline origin and matching functional repertoire do not have to coincide with a membership of the principal saline-associated clade. The repeated grouping of the sister cell pairs nevertheless provides an example of a relationship retained across draws. The substantial topological differences from random-draw concatenation of marker genes beyond the saline/freshwater split demonstrate the sensitivity to copy selection for inferring phylogenetic relationships within *Achromatium*. Consequently, a single concatenated tree cannot capture the full range of relationships contained in just a few cells.

Combining microscopy, inclusion chemistry and comparative single-cell genomics, we characterize a tropical mangrove *Achromatium* population in which most cells presented less intracellular diversity than freshwater assemblies from Lake Stechlin. Two cells, however, displayed extensive intracellular genomic diversity. Multigene phylogenies and functional gene repertoires distinguish the Guadeloupe population from freshwater references. Broader independent sampling and genomic linkage data are needed to resolve population boundaries and test the processes underlying this differentiation. Our findings demonstrate that extensive genomic diversity within individual cells can coexist with differentiation among Achromatium populations in both sequence relationships and functional potential.

## Acknowledgements

We are thankful to Janey Lee and Danielle Goudeau for technical support on this project. The work ***(proposal: 10.46936/10.25585/60001074)*** conducted by the U.S. Department of Energy Joint Genome Institute (https://ror.org/04xm1d337), a DOE Office of Science User Facility, is supported by the Office of Science of the U.S. Department of Energy operated under Contract No. DE-AC02-05CH11231.

## Author contributions

**Pierre H. Remy:** investigation (field sampling, microscopy), data curation, (ANI analyses), visualization, conceptualization, writing original draft, formal analysis, data curation. **Danny Ionescu:** conceptualization, genomic analysis, investigation, writing original draft, review and editing, data curation. **Olivier Gros:** field sampling, conceptualization, investigation, funding acquisition, writing original draft, review and editing, supervision. **Sophie-Luise Heidig:** formal analysis, visualization, bioinformatic analysis (phylogenies), writing. **George Schaible:** investigation (Raman methodology). **Tanja Woyke:** investigation, funding acquisition, review and editing. **Jean-Marie Volland:** supervision, conceptualization, project administration, review and editing.

## Data availability

Genome assemblies, functional annotation tables, computational workflow files, plotting scripts and associated outputs supporting this study are publicly available in Zenodo (version v1) at https://doi.org/10.5281/zenodo.23576787.

