## Supplementary Material. for "Single-cell genomics reveals genomic and functional differentiation between freshwater and saline *Achromatium* populations"

### Supplementary figures

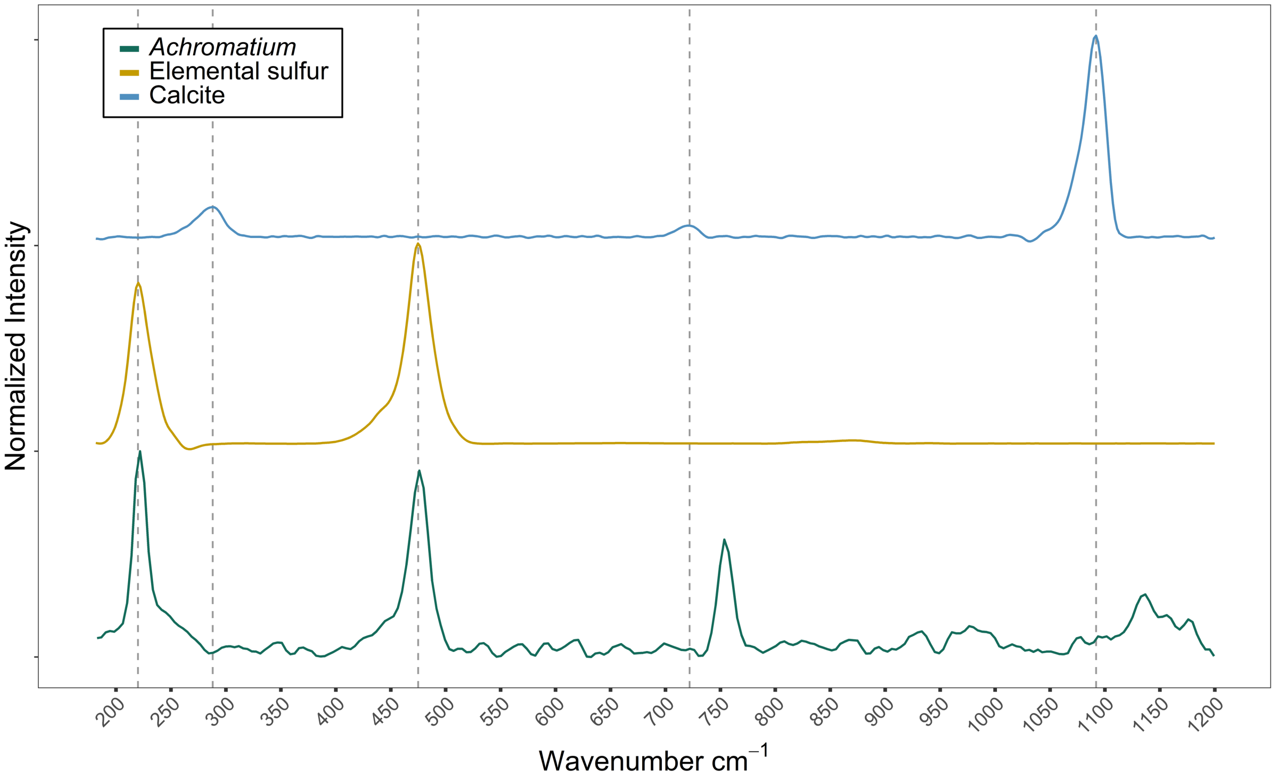


**Figure S1.** Raman spectrum of a single *Achromatium* cell compared with elemental sulfur and calcite reference spectra. The vertical lines at 222 and 476 $cm^{-1}$ match the peaks of the elemental sulfur reference. The additional peak at 754 $cm^{-1}$ was assigned to tryptophan. The *Achromatium* spectrum does not match the characteristic calcite peaks at 290 and 1092 $cm^{-1}$. The mineralogical identity of calcium-rich inclusions remained unresolved.

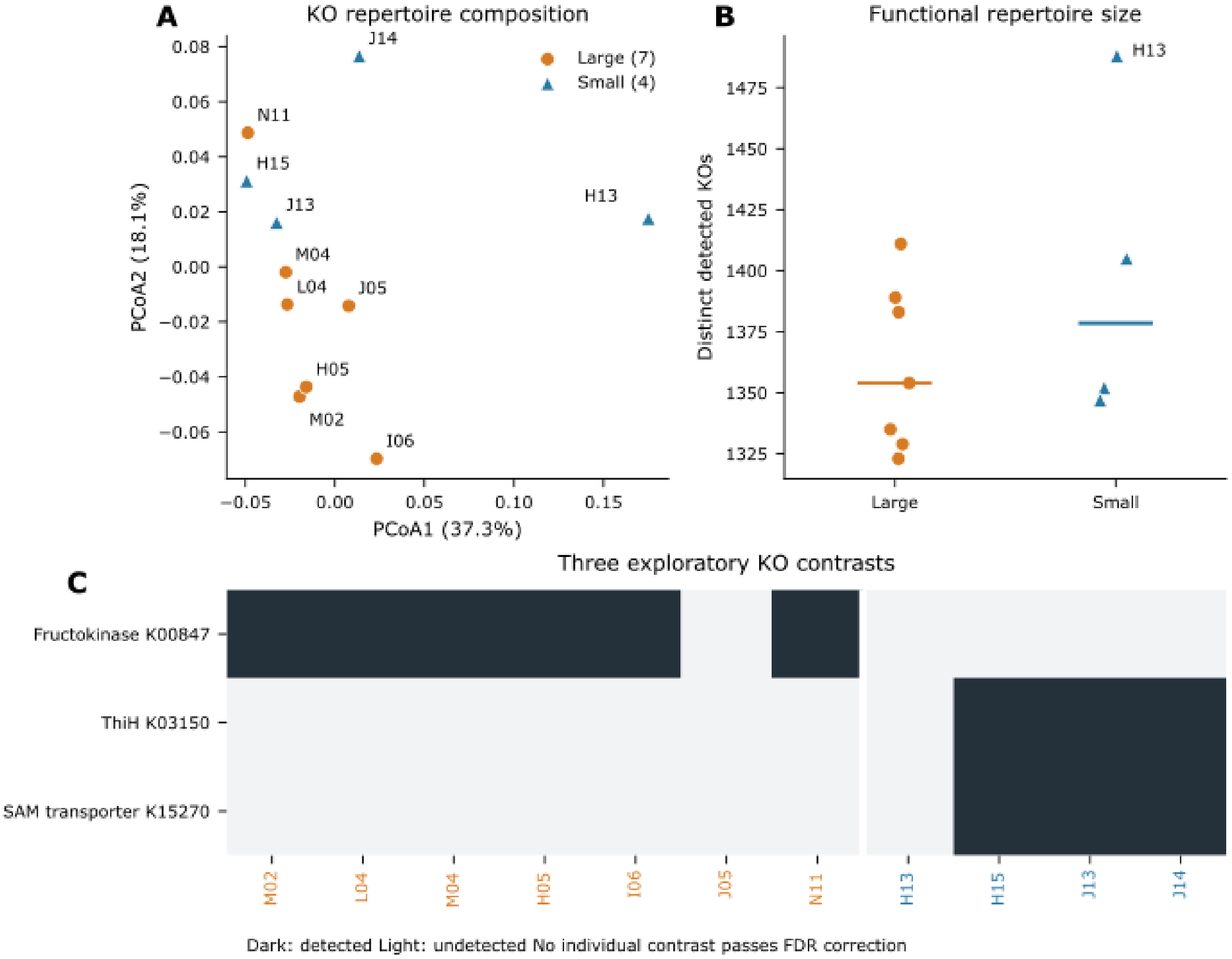


**Figure S2.** Functional comparison of large and small Guadeloupe *Achromatium* cells. Orange circles indicate seven large cells and blue triangles indicate four small cells. (A) Principal coordinates analysis of binary KO profiles using Jaccard dissimilarity. Axis percentages refer to positive PCoA inertia. (B) Distinct KO recovery per cell; horizontal lines show group medians. (C) Three exploratory KO contrasts: fructokinase K00847, ThiH K03150 and S-adenosylmethionine transporter K15270. Dark cells indicate detection and light cells indicate no detection. None of these contrasts remained significant after feature-level false-discovery-rate correction.

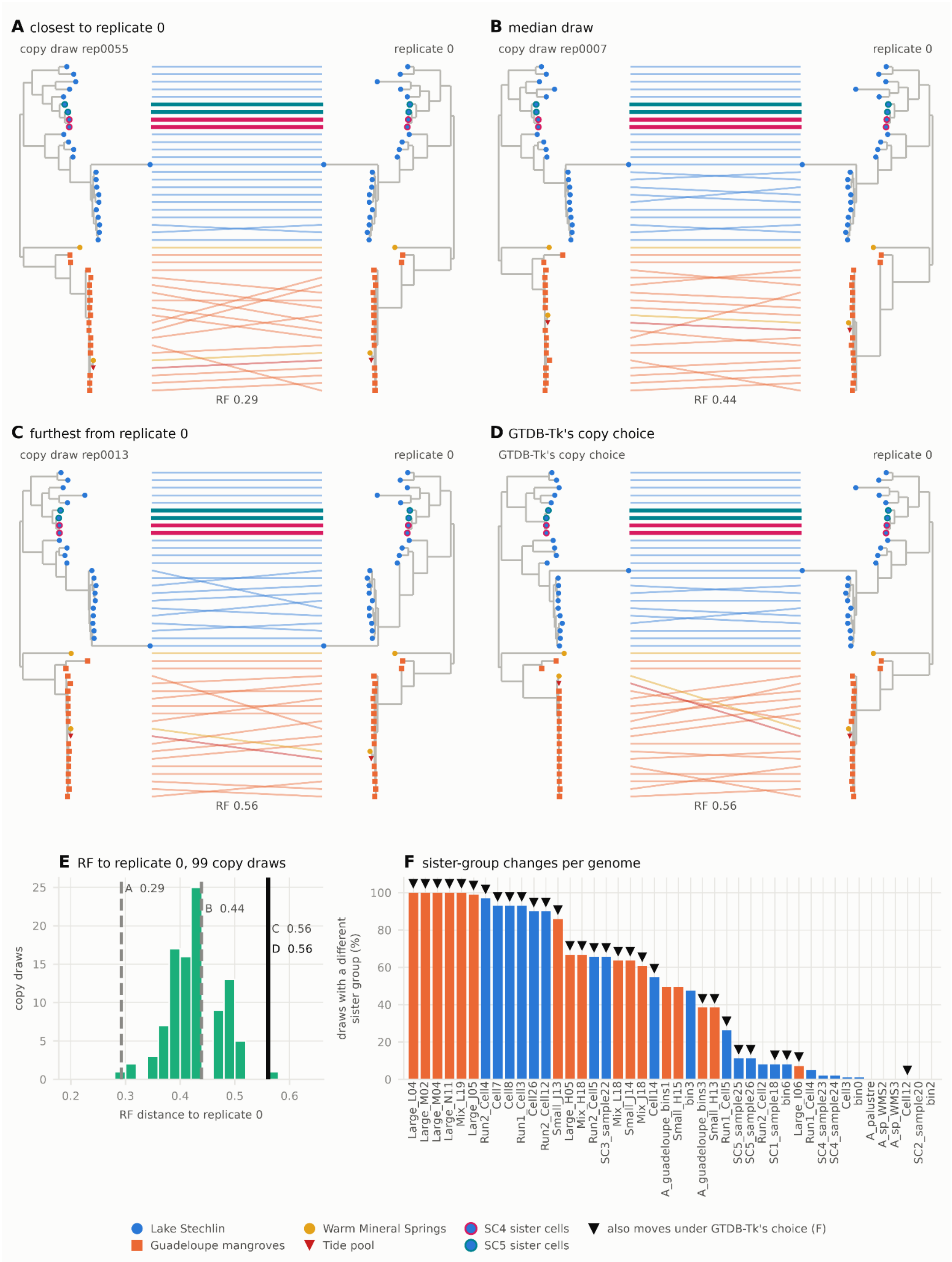


**Figure S3.** Effect of marker copy selection on concatenated *Achromatium* phylogenies. Colours identify sampling origin: blue, Lake Stechlin; orange, Guadeloupe mangroves; yellow, Warm Mineral Springs; and red, the tide-pool reference. Tip symbols are as in Fig. 6. One copy per marker per assembly was selected independently at random from the 96 retained marker families in the 44 filtered assemblies to produce 100 replicate concatenations. (A–D) Tanglegrams compare replicate 0, shown on the right, with alternative phylogenies on the left. Lines connect the same assembly between paired trees. Pink and teal rings identify the SC4 and SC5 sister-cell pairs, respectively; the four cells formed a single clade in every draw. (A–C) Comparisons with the closest, median and furthest of the other 99 random-copy phylogenies, corresponding to normalized Robinson–Foulds (RF) distances of 0.29, 0.44 and 0.56, respectively. (D) Comparison with the phylogeny obtained using GTDB-Tk's copy selections, aligned and concatenated with the same procedure as the random-copy dataset; RF = 0.56. (E) Distribution of RF distances to replicate 0 across the 99 alternative random draws; A–D identify the displayed comparisons. (F) Percentage of alternative draws in which each assembly had a different sister group from that in replicate 0. Black triangles identify assemblies that also changed sister group under GTDB-Tk's copy selection. Seven assemblies changed in at least 95% of alternative draws, including six from Guadeloupe; six assemblies never changed.

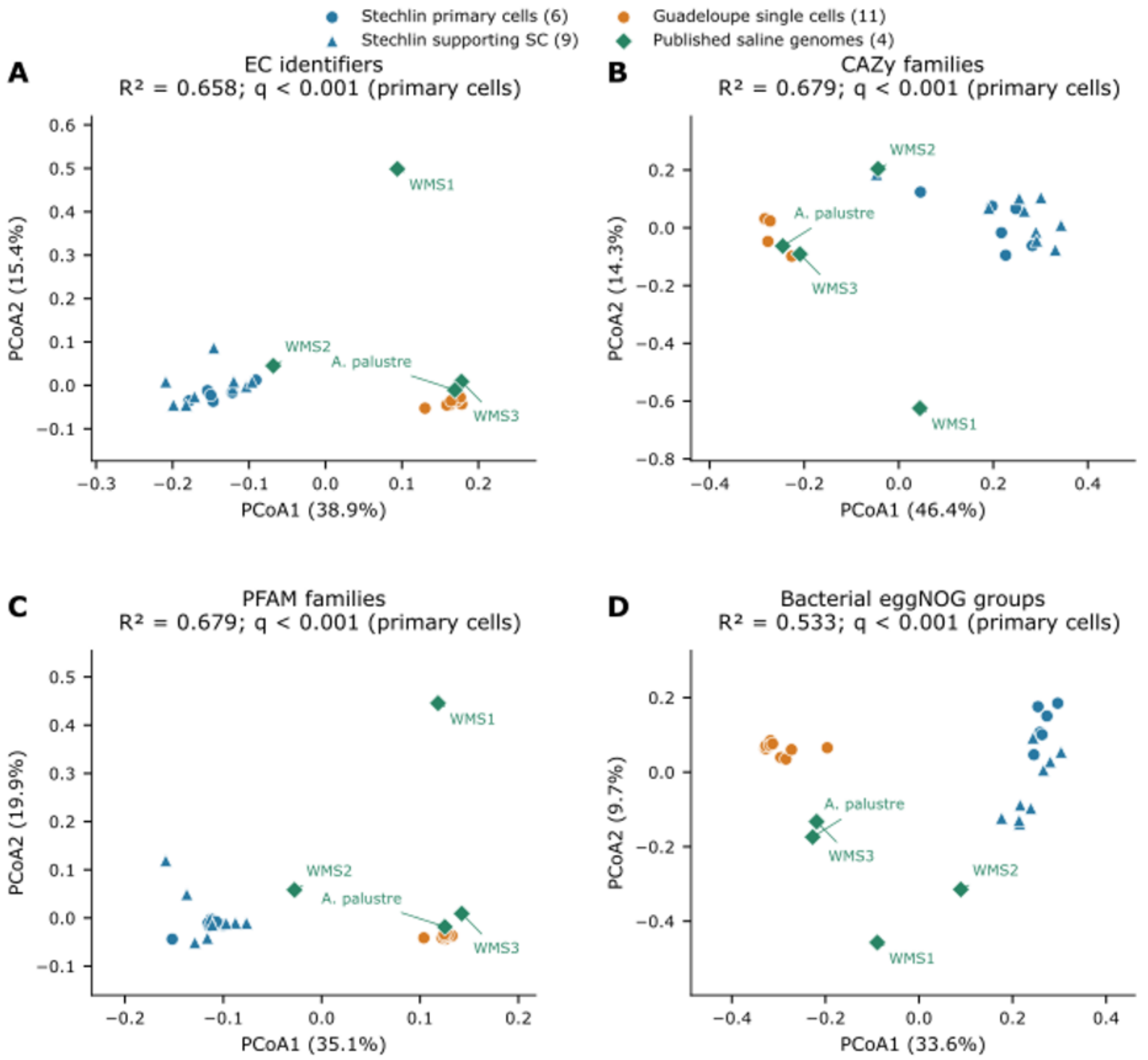


**Figure S4.** Principal coordinates analyses of *Achromatium* functional profiles across populations using alternative annotation systems. (A) Enzyme Commission (EC) identifiers. (B) CAZy families. (C) PFAM families. (D) Bacterial eggNOG groups. Ordinations use Jaccard dissimilarities calculated from binary presence–absence profiles. Blue circles indicate six primary Stechlin single-cell assemblies; blue triangles, nine supporting Stechlin split-cell (SC) assemblies; orange circles, 11 Guadeloupe single-cell assemblies; and green diamonds, four published saline genomes. Axis percentages refer to positive PCoA inertia. The PERMANOVA $R^2$ and adjusted q values above each panel refer to the primary comparison between the six Stechlin and 11 Guadeloupe single cells. Supporting SC assemblies and published saline genomes are displayed for context. The q values were adjusted across the five annotation-type comparisons.

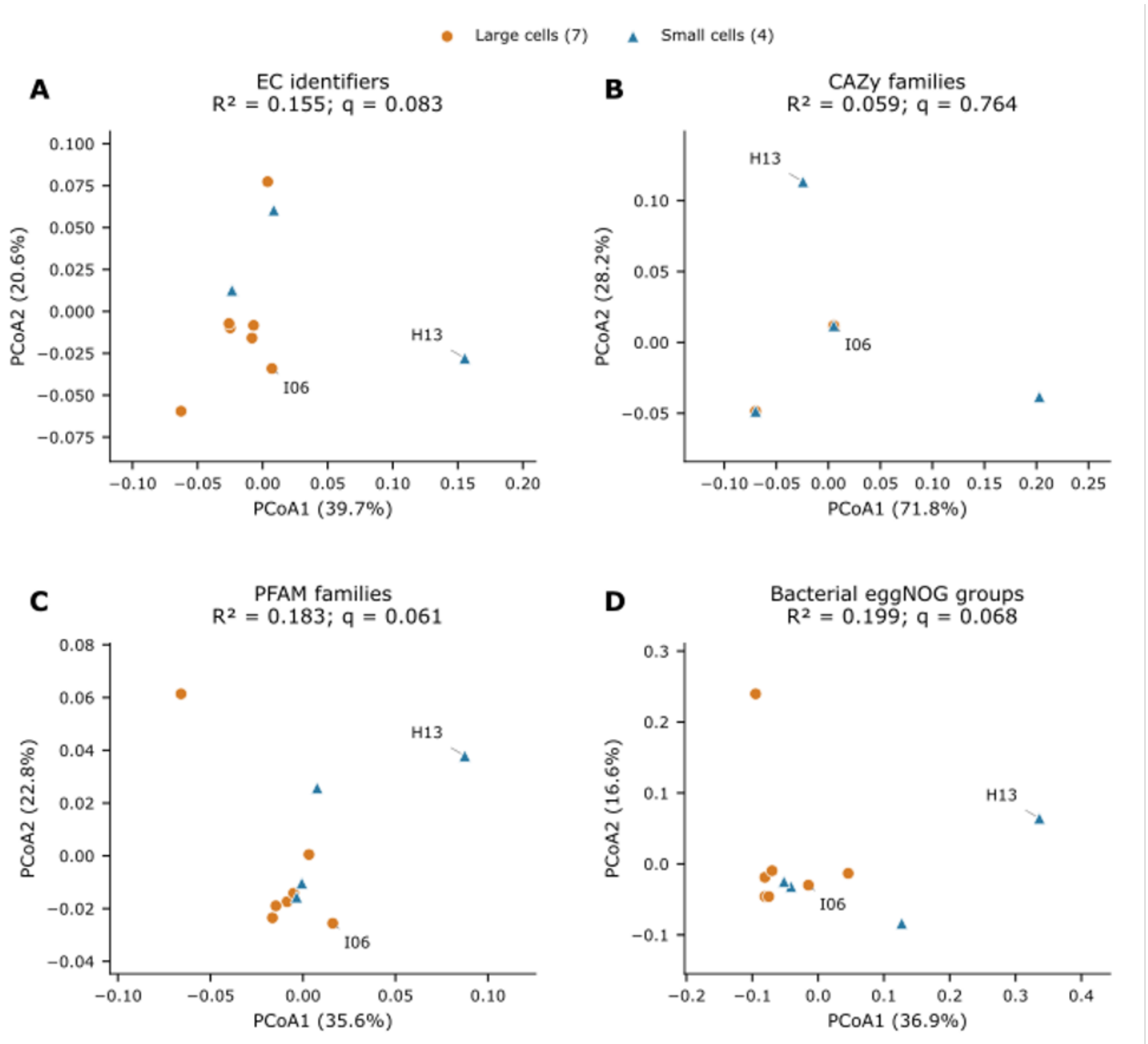


**Figure S5.** Principal coordinates analyses of functional profiles in large and small Guadeloupe *Achromatium* cells using alternative annotation systems. (A) Enzyme Commission (EC) identifiers. (B) CAZy families. (C) PFAM families. (D) Bacterial eggNOG groups. Ordinations use Jaccard dissimilarities calculated from binary presence–absence profiles. Orange circles indicate seven large cells and blue triangles indicate four small cells. Labels identify selected individual cells. Axis percentages refer to positive PCoA inertia. Panel headings report PERMANOVA $R^2$ and q values adjusted across the five annotation-type comparisons. None of the four comparisons shown was significant at $q < 0.05$.

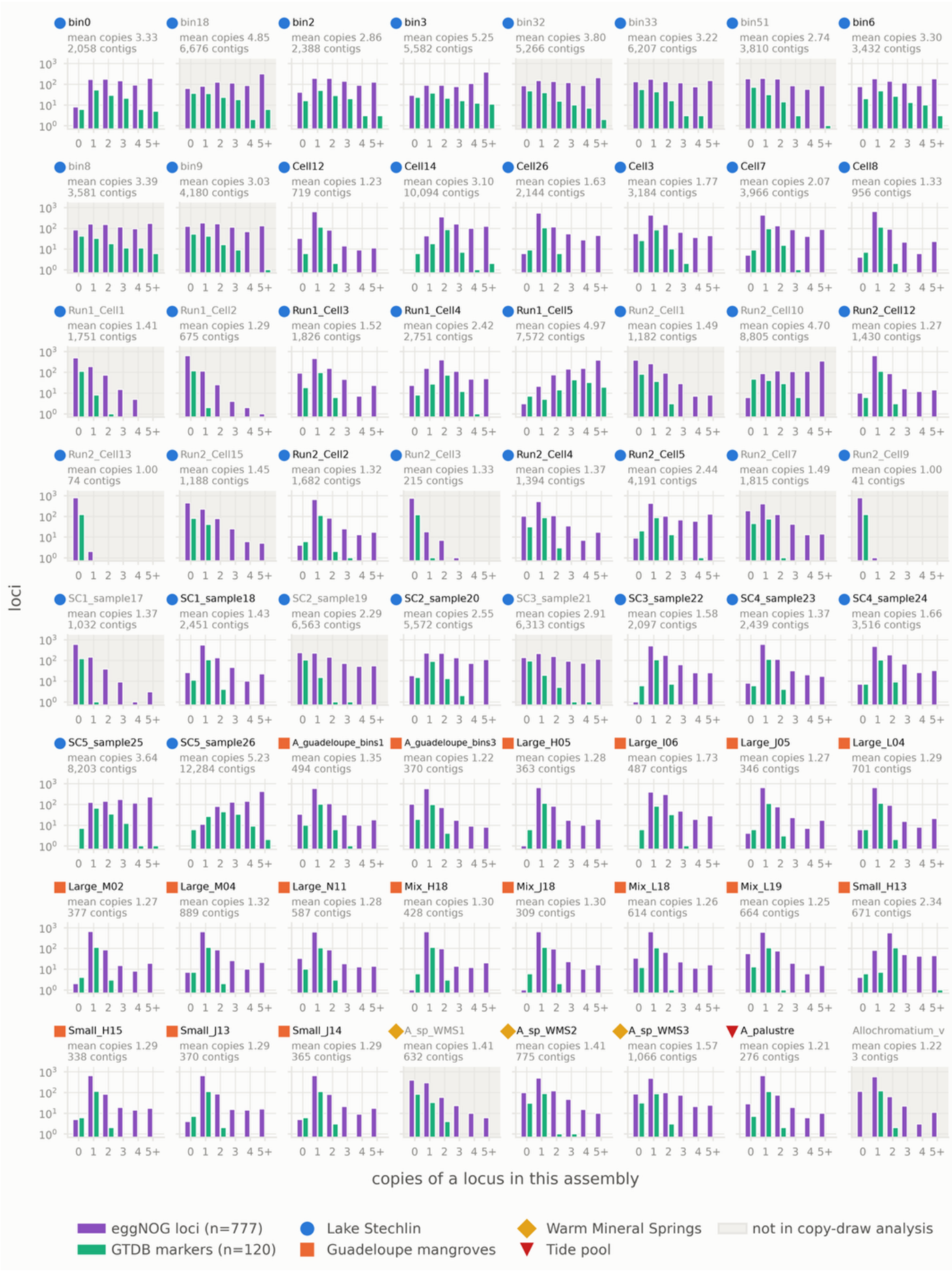


**Figure S6.** Copy-number distributions of common loci across *Achromatium* and reference assemblies. Each panel represents one assembly. Purple bars show eggNOG loci (n = 777) and green bars show GTDB bac120 markers (n = 120), grouped by the number of recovered copies per locus: 0 (missing), 1-4 or at least 5. The vertical axes show the number of loci on a logarithmic scale. Panel annotations give the reported mean copy number and contig count for each assembly. Symbols identify sampling origin: blue circles, Lake Stechlin; orange squares, Guadeloupe mangroves; yellow diamonds, Warm Mineral Springs; and red inverted triangles, the tide-pool reference. Grey panel backgrounds identify assemblies excluded from the copy-draw analysis. *Allochromatium vinosum* is included as an outgroup.

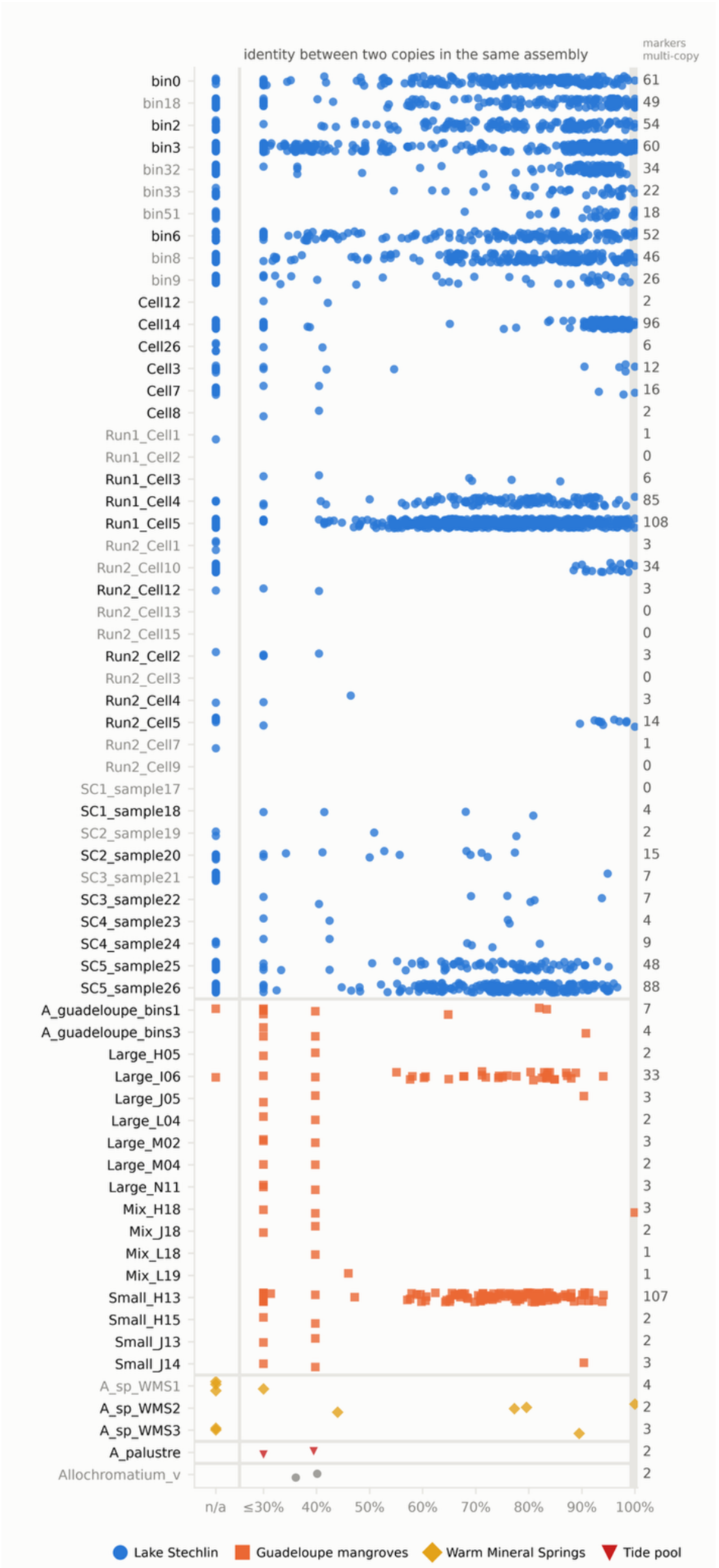


**Figure S7.** Within-assembly sequence identity between copies of GTDB bac120 markers. Each row represents one assembly, and each symbol represents a pairwise comparison. The numbers at right indicate the number of multicopy marker families per assembly. Comparisons without sufficient overlap (<30% median marker family length) between copies are displayed as n/a. Blue circles indicate Lake Stechlin assemblies, orange squares Guadeloupe mangrove assemblies, yellow diamonds Warm Mineral Springs references, and red inverted triangles the tide-pool reference. Grey symbols represent the *Allochromatium vinosum* outgroup. Grey labels identify assemblies excluded from the copy-draw analysis.

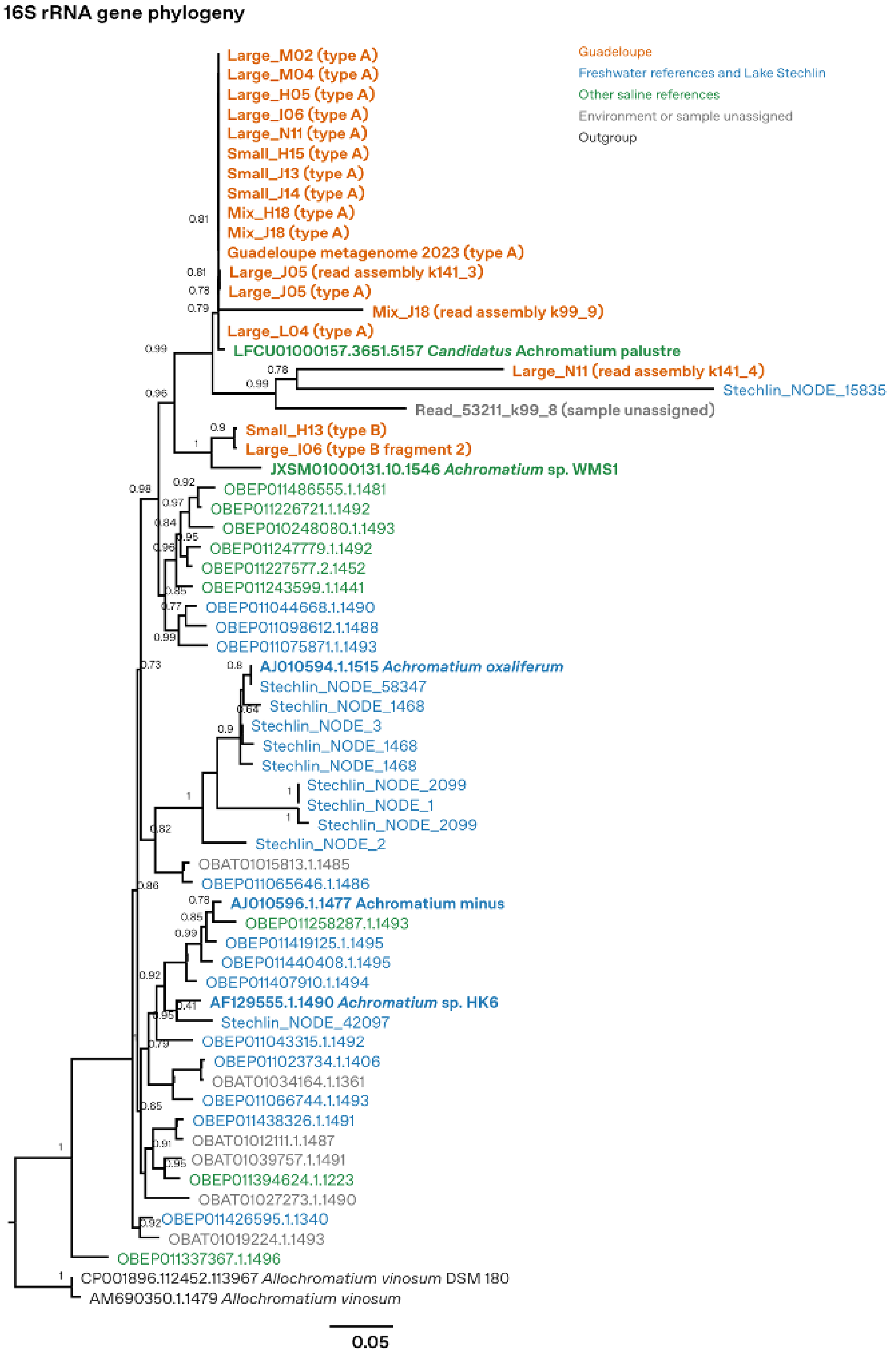


**Figure S8.** Expanded 16S rRNA gene phylogeny of *Achromatium*. Sequences from this study were analysed with selected references from SILVA release 144 and Karst et al. [41], and Lake Stechlin metagenome sequences [17]. Sequences were aligned with SINA 1.7.2 and the tree was inferred with FastTree 2 under the GTR model and rooted with *Allochromatium vinosum*. Node numbers are Shimodaira–Hasegawa-like local support values (0–1). Orange, Guadeloupe; blue, freshwater references and Lake Stechlin; green, other saline references; grey, unassigned environment or sample; black, outgroup. "Read assembly" labels denote reassemblies of sample-associated reads and do not establish sequence origin in the isolated cell. Read_53211_k99_8 remains unassigned. The scale bar represents 0.05 substitutions per site.

**Table S1.** Kraken 2 classification of the assemblies.

| Assembly | Contig count | Unclassified | Root | Bacteria/ Archaea | *Achromatium* contigs |
|---|---|---|---|---|---|
| Large_N11 | 1458 | 20 | 35 | 1403 | 587 |
| Small_H13 | 2502 | 22 | 37 | 2443 | 671 |
| Small_J14 | 1462 | 39 | 53 | 1370 | 365 |
| Mix_H18 | 2288 | 54 | 67 | 2167 | 428 |
| Mix_J18 | 841 | 21 | 21 | 799 | 309 |
| Mix_L18 | 1771 | 21 | 42 | 1708 | 614 |
| Mix_L19 | 1190 | 24 | 23 | 1143 | 664 |
| Small_H15 | 1848 | 34 | 42 | 1772 | 338 |
| Small_J13 | 1283 | 18 | 36 | 1229 | 370 |
| Large_H05 | 407 | 1 | 1 | 405 | 363 |
| Large_I06 | 926 | 7 | 9 | 910 | 487 |
| Large_J05 | 640 | 6 | 11 | 623 | 346 |
| Large_L04 (contigs) | 37220 | 18431 | 318 | 18471 | 1950 |
| Large_L04 (scaffolds) | 35885 | 17587 | 317 | 17981 | 1933 |
| Large_M02 | 600 | 5 | 6 | 589 | 377 |
| Large_M04 (contigs) | 51975 | 24913 | 470 | 26592 | 2751 |
| Large_M04 (scaffolds) | 51705 | 24776 | 469 | 26460 | 2718 |
| NoTemTmaFil12C2 | 2715 | 32 | 7 | 2676 | 0 |
| NoTemTmaFil12D1 | 2351 | 35 | 16 | 2300 | 0 |
| NoTemTmaFil12E1 | 2076 | 15 | 13 | 2048 | 0 |
| NoTemTmaFil12J2 | 2417 | 23 | 6 | 2388 | 0 |
| NoTemTmaFil12L2 | 2358 | 25 | 4 | 2329 | 0 |
